# Generative machine learning produces kinetic models that accurately characterize intracellular metabolic states

**DOI:** 10.1101/2023.02.21.529387

**Authors:** Subham Choudhury, Bharath Narayanan, Michael Moret, Vassily Hatzimanikatis, Ljubisa Miskovic

**Affiliations:** Laboratory of Computational Systems Biology (LCSB), Ecole Polytechnique Fédérale de Lausanne (EPFL), Lausanne, CH-1015, Switzerland; Department of Oncology, University of Cambridge, Cambridge, CB2 0XZ, UK; Department of Genetics, Harvard Medical School, Boston, MA 02115, USA

**Author notes:** **Correspondence:** Ljubisa Miskovic, Laboratory of Computational Systems Biotechnology (LCSB), École Polytechnique Fédérale de Lausanne (EPFL), CH-1015 Lausanne, Switzerland; Vassily Hatzimanikatis, Laboratory of Computational Systems Biotechnology (LCSB), École Polytechnique Fédérale de Lausanne (EPFL), CH-1015 Lausanne, Switzerland.

**Keywords:** metabolism, omics data integration, large-scale and genome-scale kinetic models, machine learning, evolution strategies, *E. coli*, kinetic parameters, nonlinear dynamics

## Abstract

Generating large omics datasets has become routine practice to gain insights into cellular processes, yet deciphering such massive datasets and determining intracellular metabolic states remains challenging. Kinetic models of metabolism play a critical role in integrating omics data, as they provide explicit connections between metabolite concentrations, metabolic fluxes, and enzyme levels. Nevertheless, the challenges associated with determining the kinetic parameters that underlie cellular physiology pose significant obstacles to the broader acceptance and adoption of these models within the research community. Here, we present RENAISSANCE, a generative machine learning framework for efficiently parameterizing large-scale kinetic models with dynamic properties matching experimental observations. Through seamless integration and consolidation of diverse omics data and other relevant information, like extracellular medium composition, physicochemical data, and expertise of domain specialists, we show that the proposed framework accurately characterizes unknown intracellular metabolic states, including metabolic fluxes and metabolite concentrations in *E. coli*’s metabolic network. Moreover, we show that RENAISSANCE successfully estimates missing kinetic parameters and reconciles them with sparse and noisy experimental data, resulting in a substantial reduction in parameter uncertainty and a notable improvement in the accuracy and reliability of the parameter estimates. The proposed framework will be invaluable for researchers who seek to analyze metabolic variations involving changes in metabolite and enzyme levels and enzyme activity in health and biotechnological studies.

---

Advancement in biotechnology and health sciences hinges heavily on our capability to integrate different varieties of data produced by high-throughput techniques and obtain coherent insights into cellular processes^1–3^. Considerable effort has been invested in using genome-scale models, mathematical representations of metabolic information about living organisms, to reconcile and make sense of such constantly growing disparate datasets^4,5^. Genome-scale models integrate omics data by considering constraints imposed by genetics and physicochemical laws^6–10^. For instance, researchers use inequality constraints stemming from the second law of thermodynamics to relate metabolic fluxes (fluxome) to metabolite profiles (metabolome)^11–14^. However, data integration using such inequality constraints results in significant uncertainty about intracellular metabolic states^15^. Consequently, despite the availability of large omics datasets, determining the exact intracellular levels of metabolite profiles and metabolic reaction rates with these constraint-based models remains elusive.

Kinetic models of metabolism can address these issues by consolidating several types of omics data, such as metabolomics, fluxomics, transcriptomics, and proteomics, within a common and coherent mathematical framework^16^. Indeed, these models contain information about enzyme kinetics and metabolic regulation, allowing them to explicitly couple metabolite concentrations, metabolic reaction rates, and enzyme levels through mechanistic relations. Additionally, unlike constraint-based models, kinetic models capture time-dependent responses of cellular metabolism. Taken altogether, these models show great promise for addressing complex phenomena in biomedical sciences and biotechnology, such as metabolic reprogramming in the tumor microenvironment and disease^17–19^, relationships between cancer, metabolism, and circadian rhythms^20^, dynamics of drug absorption and drug metabolism^21^, and engineering and modulating cell phenotypes^22–24^.

Despite the capacity of kinetic models to reconcile data and identify metabolic features associated with phenotype, the application of these models is somewhat limited^16,25–30^. The major challenge in developing kinetic models is the lack of knowledge about the characteristic kinetic parameter values that govern the cellular physiology of the studied organism *in vivo*. Overcoming this requires employing intricate computational procedures and the extensive expertise of researchers. It is often impractical to build and use these models for studying multiple physiological conditions and large cohorts^31^. Therefore, there is a need for accelerated approaches for parameterizing kinetic models that would allow the broader research community access to these models.

Recent efforts employing new tailor-made parameterization^28^ and machine learning^32–34^ improved the efficiency of constructing near-genome-scale kinetic models. Nevertheless, challenges remain regarding extensive computational time^28^ and the need for training data from traditional kinetic modeling approaches^32–34^. Here, we present RENAISSANCE (REconstruction of dyNAmIc models through Stratified Sampling using Artificial Neural networks and Concepts of Evolution strategies), a machine learning framework that efficiently parameterizes biologically relevant kinetic models of metabolism without requiring training data. The behavior of parameterized kinetic models is highly nonlinear yet deterministic and depends on the intracellular state, defined by network topology and integrated data. To capture this nonlinear behavior, we use feed-forward neural networks of comparable complexity and optimize them with Natural Evolution Strategies (NES)^35,36^ to obtain kinetic models with desired properties (Figure 1a). This dramatically reduces the extensive computation time required by traditional kinetic modeling methods, thus allowing its broad utilization for high-throughput dynamical studies of metabolism. We showcase RENAISSANCE through three studies: (i) generating a population of large-scale dynamic models of *E. coli* metabolism, (ii) characterizing intracellular metabolic states in the *E. coli* metabolic network accurately, and (iii) integrating and reconciling available experimental kinetic data.

**Figure 1.**
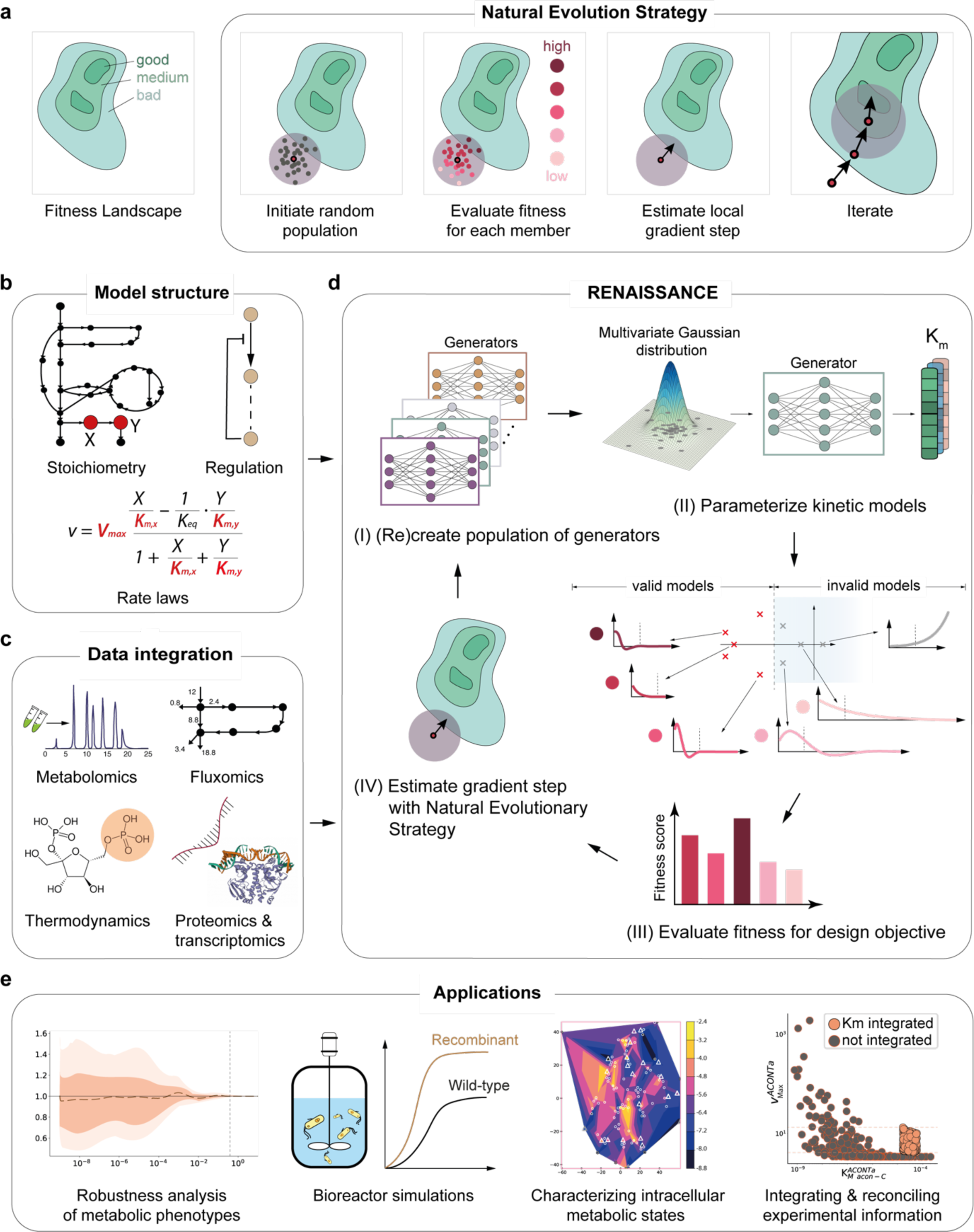
Overview and applications of the RENAISSANCE framework. **a**, A Natural Evolutionary Strategy (NES) algorithm iteratively generates candidate solutions for an optimization problem based on their assigned fitness scores until satisfactory solutions are obtained. **b**, Context-specific structural properties of the metabolic networks are established and incorporated into the model. **c**, Once the model structure is fixed, available omics data are integrated into the model. **d**, Generators for parameterizing biologically relevant (valid) kinetic models are optimized iteratively in four steps to meet the design objective: a population of generators is randomly initialized (step I); generators produce parameters needed to parameterize kinetic models (step II); the fitness of the kinetic models (circles and bars in shades of red) is assessed based on the largest eigenvalues of the Jacobian (red and grey crosses) corresponding to the dominant time constants of the model responses (Methods); the generator is assigned a score based on this performance (step III); the rewards for each generator are fed back to NES to find the best-performing generator (step IV); the best-performing generator is then perturbed to obtain the next generation of generators (step I). **e**, A few applications of RENAISSANCE-generated models presented in this paper.

## Results

### Devising a framework for parameterizing biologically relevant kinetic models

We developed RENAISSANCE, a machine-learning framework for parameterizing biologically relevant kinetic models. These models are consistent with experimentally observed steady states and produce dynamic metabolic responses with timescales^37^ that match experimental observations in cellular organisms. The input to RENAISSANCE is a steady-state profile of metabolite concentrations and metabolic fluxes computed by integrating structural properties of the metabolic network (stoichiometry, regulatory structure, rate laws) and available data (metabolomics, fluxomics, thermodynamics, proteomics, and transcriptomics) into the model (Figure 1b, c, Methods).

RENAISSANCE uses feed-forward neural networks (generators) to parameterize kinetic models, with the size of generator networks dictated by the complexity of the kinetic model. Using NES, it optimizes the weights of generators in four iterative steps until they produce biologically relevant models (Figure 1d, Methods). The iterative process starts by initializing a population of generators with random weights (step I). The use of multiple generators facilitates a more thorough and more efficient exploration of parametric space. Each generator takes multivariate Gaussian noise as input and produces a batch of kinetic parameters consistent with the network structure and integrated data. These parameter sets are then used to parameterize the kinetic model (step II). Next, we evaluate the dynamics of each parameterized model by computing the eigenvalues of its Jacobian and the corresponding dominant time constants (Methods). These quantities allow us to assess if the generated kinetic models have dynamic responses corresponding to experimental observations (valid models) or not (invalid models). Based on this evaluation, we assign a reward to the generator (step III). NES repeats steps II and III for every generator in the population, followed by normalizing all rewards. The weights of the ‘parent’ generator for the next generation are then obtained by using the weights of all the members of the previous generation, weighted by their normalized rewards. Although high-performing generators have a greater impact on the weight of the parent generator in the next generation, lower-performing individuals also contribute. NES subsequently mutates this ‘parent’ generator by injecting a predefined noise level into its weights, thus recreating a population of generators (Step I). We iterate steps I-IV until we obtain a generator that meets the user-defined design objective, such as maximizing the incidence of biologically relevant kinetic models (Methods).

The generated kinetic models are versatile and applicable to a broad range of metabolism studies (Figure 1e).

### Generating large-scale kinetic models of *E. coli* metabolism

We studied the anthranilate-producing *E. coli* strain, W3110 *trpD9923*, to test and validate RENAISSANCE. The kinetic model structure for this strain, adopted from Narayanan et al.^38^ consisted of 113 nonlinear ordinary differential equations (ODEs) parameterized by 502 kinetic parameters, including 384 Michaelis constants, *K*_*M*_s (Methods, Supplementary Figure 4). It encompasses 123 reactions and describes core metabolic pathways, including glycolysis, the pentose phosphate pathway (PPP), the tricarboxylic cycle (TCA), anaplerotic reactions, the shikimate pathway, glutamine synthesis, and a lumped reaction for growth (Methods, Supplementary Figure 5). The objective was to find kinetic parameters resulting in dynamic models consistent with an experimentally observed doubling time of 134 minutes for the studied *E. coli* strain^39^. A valid kinetic model satisfying this requirement should produce metabolic responses with the dominant time constant of 24 mins, corresponding to having the largest eigenvalue *λ*_*max*_ < −2.5 (Methods).

We used thermodynamics-based flux balance analysis^13,40^ to integrate experimental data^39^ and compute 5,000 steady-state profiles of metabolite concentrations and fluxes (Methods). We selected one of these profiles as input for RENAISSANCE (Methods) and identified a set of hyperparameters yielding the best framework performance with a 3-layer generator neural network (Methods, Supplementary Notes 2-4). RENAISSANCE was then executed for 50 evolution generations using the optimized settings. We repeated the optimization process 10 times with a randomly initialized generator population to obtain statistical replicates. For each generation, we generated 100 kinetic parameter sets for every generator in the population and computed the maximum eigenvalue, *λ*_*max*_, for each parameter set. To evaluate and rank the generators, we used the incidence of valid models, defined as the proportion of the generated models that are valid (with *λ*_*max*_ < −2.5, Methods). We observed that the incidence of valid models steadily increases with the number of generations, with the mean incidence converging around 92% after 50 generations (Figure 2a, thick black line, Supplementary Figure 14, 16). For some repeats, we could achieve incidence up to 100% (Figure 2a, green-shaded region).

**Fig. 2.**
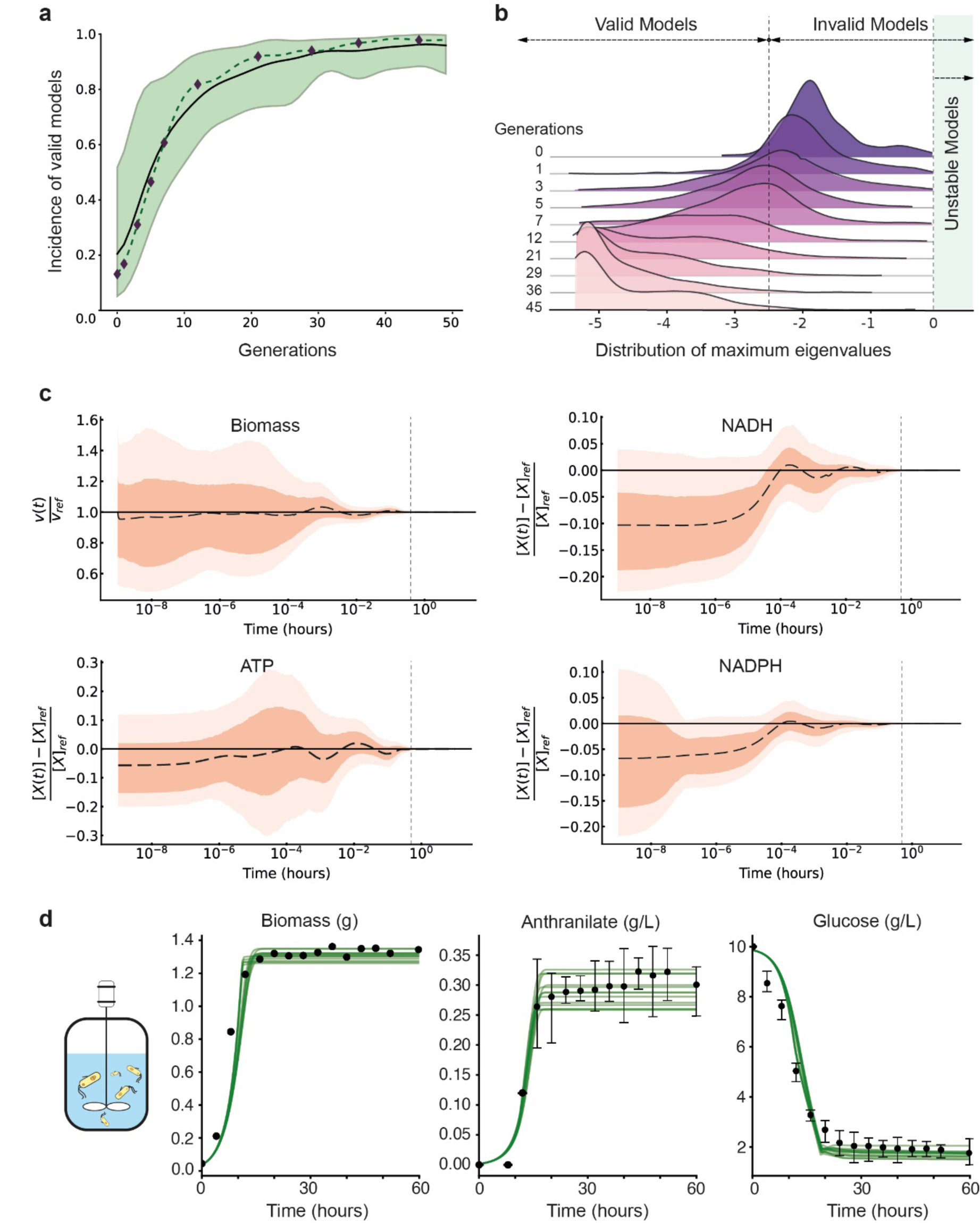
Generation, validation, and application of RENAISSANCE-parameterized kinetic models. **a**, The incidence of models exhibiting the desired dynamic properties increases with the number of generations, as indicated by the mean incidence (black line) and the maximum and minimum incidence (green-shaded region) observed over 10 statistical repeats for every generation. The dashed line indicates the incidence of a repeat with fast convergence. The black diamonds indicate the generators selected for subsequent analysis from that repeat. **b**, The distribution of the maximum eigenvalues (*λ*_*max*_) for the generated models over generations. The vertical dashed lines indicate *λ*_*max*_ = −2.5 (left) and *λ*_*max*_ = 0 (right). *Robustness analysis:* **c**, The time evolution of the normalized perturbed biomass, *v*(*t*)/*v*_*ref*_, (upper left) and concentrations, (*X*(*t*) − *X*_*ref*_)/*X*_*ref*_, Nicotinamide adenine dinucleotide reduced, NADH (upper right), Adenosine triphosphate, ATP (lower left), and Nicotinamide adenine dinucleotide phosphate reduced, NADPH (lower right), respectively: the mean response (dashed black line), the 25^th^-75^th^ percentile (dark orange region), and the 5^th^-95^th^ percentile (light orange region) of the ensemble of responses. The vertical dashed line corresponds to *t* = 24 *mins*. *Bioreactor simulations:* **d**, The time evolution of biomass (left), glucose concentration (middle), and anthranilate concentration (right) in the bioreactor runs of the 13 models closely fitting the experimental data^39^. Black dots and error bars represent the mean and standard deviation from triplicate experiments.

For further analysis of the generated models, we selected a statistical repeat with fast convergence (Figure 2a, dashed line) and chose 10 generators from that repeat with monotonically increasing incidence over generations (Figure 2a, black diamonds). For each of the 10 chosen generators, we generated 500 kinetic parameter sets and examined the distribution of the resulting maximum eigenvalues (Figure 2b). Remarkably, the generated models gradually shifted over the optimization process from having slow dynamics (*λ*_*max*_ > −2.5) to having fast dynamics, with the metabolic processes settling before the subsequent cell division, indicating that RENAISSANCE-generated models could capture the experimentally observed dynamics.

Since cellular organisms maintain phenotypic stability when faced with perturbations^41^, the generated models that describe cellular metabolism should possess the same property. To test the robustness of the models, we perturbed the steady-state metabolite concentrations up to ±50% and verified if the perturbed system returned to the steady state. For this purpose, we generated 1000 relevant kinetic models using the final of 10 selected generators (Figure 2a, generation 45), chosen for yielding the highest incidence of valid models. Inspection of the time evolution of the normalized biomass showed that the biomass returned to the reference steady state (*v*(*t*)/*v*_*ref*_ = 1) within 24 minutes for 100% of the perturbed models (Figure 2c). Similarly, the perturbed time responses of a few critical metabolites, namely, NADH, ATP, and NADPH, returned to their steady-state values within 24 minutes for 99.9%, 99.9%, and 100% of the 1000 generated kinetic models, respectively (Figure 2c). Examining every cytosolic metabolite collectively revealed that 75.4% of the models returned to the steady state within 24 minutes and 93.1% returned within 34 mins, demonstrating that the generated kinetic models are robust and obey imposed context-specific observable biophysical timescale constraints.

Next, we tested the generated models in nonlinear dynamic bioreactor simulations closely mimicking real-world experimental conditions^38,39^. The temporal evolution of biomass production showed similar trends as typical experimental observations with clear exponential and stationary phases of *E. coli* growth (Figure 2d, Supplementary Note 6, Supplementary Figure 6). Similarly, glucose uptake and anthranilate production also reproduce trends observed in experiments with glucose consumption halted and anthranilate production saturating at around 20 hours. This study indicates that the RENAISSANCE models can accurately reproduce the physiologically observable and emergent properties of cellular metabolism, even without implicit training to reproduce fermentation experiments.

### Characterizing the intracellular states of *E. coli* metabolism

Accurately determining the intracellular levels of metabolite profiles and metabolic reaction rates is crucial for associating metabolic signatures with phenotype. Yet, our capabilities to establish the intracellular metabolic state are limited. Even with the ever-increasing availability of physiological and omics data, a significant amount of uncertainty in the intracellular states remains. We propose using kinetic models to reduce this uncertainty because of their explicit coupling of enzyme levels, metabolite concentrations, and metabolic fluxes. Moreover, kinetic models allow us to consider dynamic constraints in addition to steady-state data, thus allowing us further uncertainty reduction.

After integrating available physiology and omics data^42,43,39,44^ using the constraint-based thermodynamics-based flux balance analysis^40^, significant uncertainty was present in the intracellular metabolic state as indicated by the wide ranges of metabolite concentrations and metabolic fluxes. We sampled 5000 steady-state profiles of metabolite concentrations and metabolic fluxes from this uncertain space and deployed RENAISSANCE to find the fastest possible dynamics (maximum negative eigenvalues, *λ*_*max*_) for each steady state (Methods, Supplementary figure 7). We visualized the steady-state profiles by performing dimension reduction with Principal Component Analysis (PCA)^45^ and t-distributed Stochastic Neighbor Embedding (t-SNE)^46^ (Methods) and colored each steady-state profile according to the obtained *λ*_*max*_ (Figure 3a). We observed a high variation in the dynamics (*λ*_*max*_) of the studied steady-state profiles (Figure 3c, blue distribution). Of 5000 steady-state profiles, 918 (18.4%) had *λ*_*max*_ larger than −2.5, meaning these intracellular metabolic states could not correspond to the experimental observations. Indeed, the dynamic responses corresponding to these states have a time constant superior to 24 mins, i.e., slower than the experimental observations.

**Fig. 3.**
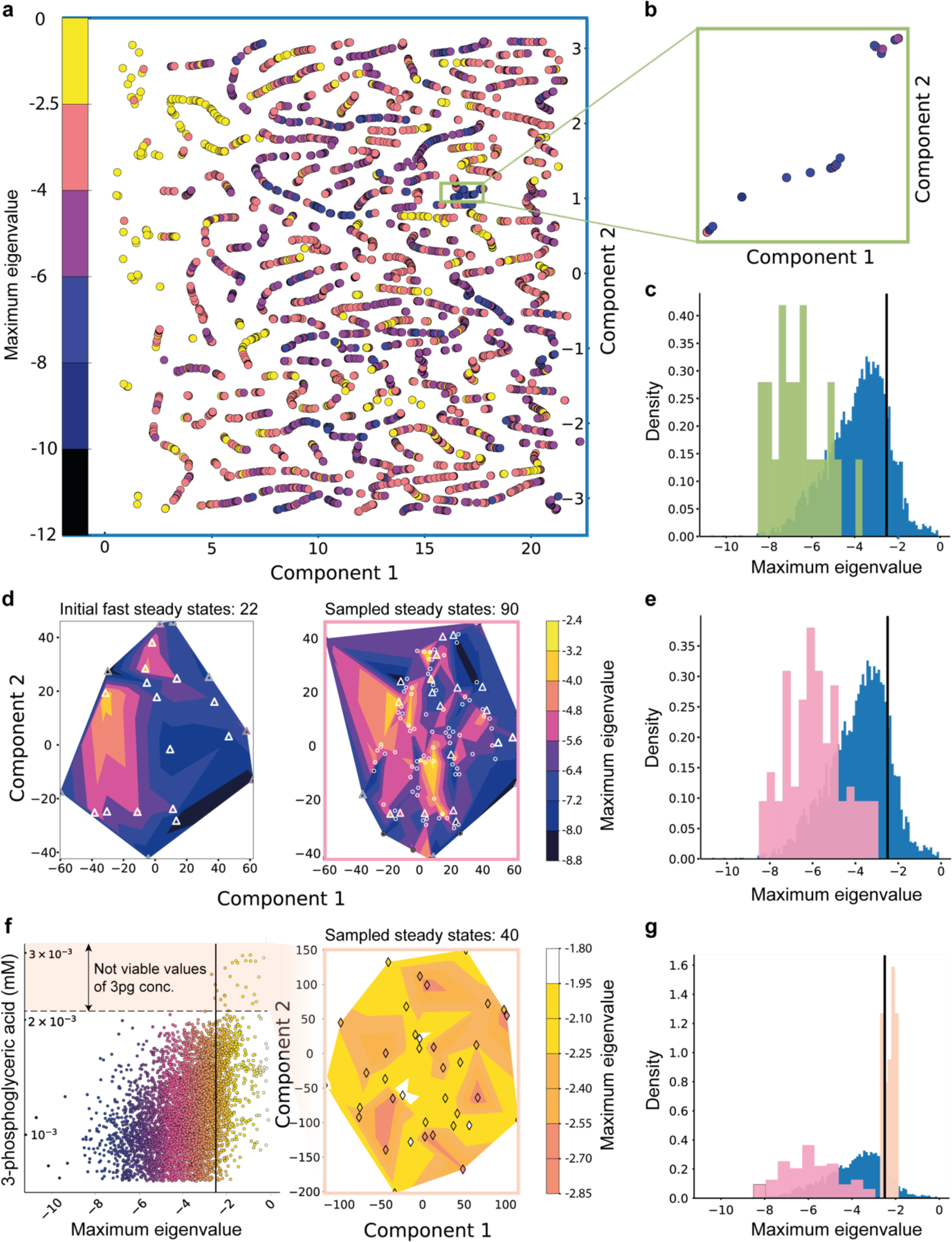
Dynamic characterization reduces uncertainty in intracellular metabolic states. **a**, The two-dimensional representation of the fastest linearized dynamic modes (corresponding to the maximum eigenvalue *λ*_*max*_) of 5000 intracellular steady states (reaction fluxes and metabolite concentrations) obtained with Principal Component Analysis (PCA) and t-Distributed Stochastic Neighbor Embedding (t-SNE)^46^ (Methods). Each point represents a steady state, with its color indicating the corresponding *λ*_*max*_ value computed by RENAISSANCE. **b**, Magnified view of 22 neighboring steady states with fast dynamics (−3.8 ≤ *λ*_*max*_ ≤ −8.5). The color scheme is the same as in 3a. **c**, Distributions of the fastest linearized dynamic (*λ*_*max*_) for all 5000 steady states (blue) and the 22 steady states shown in 3b (green). **d**, Left: the linearized dynamics landscape of the 22 fast steady states in the reduced space (Methods). The triangles represent the location of the steady states in the landscape. Right: The landscape on the left is enhanced by sampling 90 additional steady states in the neighborhood of the initial 22 steady states. The circles represent the location of the newly sampled steady states in the same landscape as on the left. **e**, Distributions of the fastest linearized dynamic (*λ*_*max*_) for all steady states (blue) and 90 steady states sampled in 3d (pink). **f**, Left: concentration of 3-phosphoglyceric acid (mM) vs. the fastest linearized dynamic in every steady state. The color scheme is the same as that in 3a. The horizontal black line indicates the cutoff for valid models (*λ*_*max*_ = −2.5). The peach-shaded region shows the range of 3-phosphoglyceric acid concentration that does not allow fast dynamics. Right: the dynamic landscape of 40 steady states sampled by constraining the metabolite concentrations of 30 metabolites to ranges that do not support fast dynamics. The diamonds represent the location of the steady states. **g**, Distributions of the fastest linearized dynamic (*λ*_*max*_) for the 40 steady states sampled in 3f (peach), compared to all steady states (blue) and those sampled in 3d (pink).

As t-SNE optimizes the preservation of local distances between points when projecting them from a high-dimensional space to a lower-dimensional one^46,47^, we hypothesized that sampling from a region containing closely positioned steady-state profiles associated with fast dynamics (Figure 3a, blue dots) would yield steady-state profiles that satisfy dynamic requirements. Conversely, sampling from a region around adjacent profiles corresponding to slow dynamics (Figure 3a, yellow dots) would likely result in profiles not meeting dynamic requirements.

To test this hypothesis, we selected one of these local regions (Figure 3b), which contained 22 steady states with fast dynamics with −3.8 ≤ *λ*_*max*_ ≤ −8.5 (Figure 3c, green distribution), and analyzed its neighborhood (Figure 3d, left). We sampled 90 additional steady states within this neighborhood from the Gaussian distribution with a mean and standard deviation estimated on the initial 22 steady states. The sampled steady states allowed us to improve the resolution of the initial dynamic landscape (Figure 3d, right, circles). Crucially, the sampled steady states had linearized dynamics in the same range as the initial 22 states (Figure 3d, e), confirming our hypothesis. Therefore, RENAISSANCE allows us to select subsets of intracellular states consistent with experimentally observed dynamics and generate additional ones with the same characteristics. Moreover, it allows us to discard subregions with experimentally inconsistent states, thus reducing uncertainty. Indeed, sampling from a region containing closely positioned steady-state profiles associated with slow dynamics yielded steady states corresponding to similarly slow dynamics (Supplementary Figure 13).

We next examined individual metabolite concentrations of the 5000 steady-state profiles to identify patterns corresponding to the experimentally observed phenotype. We observed a clear bias in the dynamics depending on the concentrations for some of the metabolites (Figure 3f, Supplementary Figure 7). For example, in the case of 3-phosphoglyceric acid (3pg), we obtain models with relevant dynamics only when the concentration of this metabolite is less than ∼ 0.002 *mM*. In contrast, steady-state profiles with 3pg concentrations between 0.002 − 0.003 *mM* do not have relevant dynamics (Figure 3f). To investigate this further, we identified 30 cytosolic metabolites that showed such concentration biases by visual inspection (Supplementary Figure 8) and sampled 40 new steady states from the same Gaussian distribution as before (Figure 3d, left) but constrained the selected 30 metabolites to concentration ranges that do not support relevant dynamics (e.g., peach shaded region in Figure 3f). As expected, almost all of these new intracellular states did not yield models with relevant dynamics (Figure 3f, right and 3g). This result demonstrates that information stemming from the dynamic responses can be used to constrain values of intracellular metabolites to specific ranges.

Overall, dynamic characterization of a broad range of intracellular states allows us to reduce uncertainty at the level of steady-state profiles, as well as individual metabolite concentrations and metabolic fluxes.

### Integration and reconciliation of experimental information

Experimentally measured Michaelis constants, *K*_*M*_s, are curated in comprehensive databases like BRENDA^48^. However, as we transition to large genome-scale kinetic models, a vast majority of the associated kinetic parameters remain unknown. Integrating experimental results from *in vivo* and *in vitro* studies, despite the disparities in their parameter values, can help further constrain uncertainty and lead to a more accurate description of intracellular metabolic states. To this end, we retrieved experimentally measured values for 108 out of 384 *K*_*M*_s in our model from BRENDA (Methods).

To investigate how the integrated kinetic data constrain unknown kinetic parameters, we started by integrating 4 *K*_*M*_ values of aconitase (ACONTa, b) from the citric acid cycle (Figure 4a, Methods), obtained generators with a high incidence of valid models (>99%), and generated 500 valid kinetic models (Supplementary Figure 9). To quantify the effect of integrating one experimental *K*_*M*_ value on the generated values of the other kinetic parameters, we compared the estimates of the other *K*_*M*_s and maximum velocities, *v*_*max*_, with ones obtained when no kinetic parameters were integrated.

**Fig. 4.**
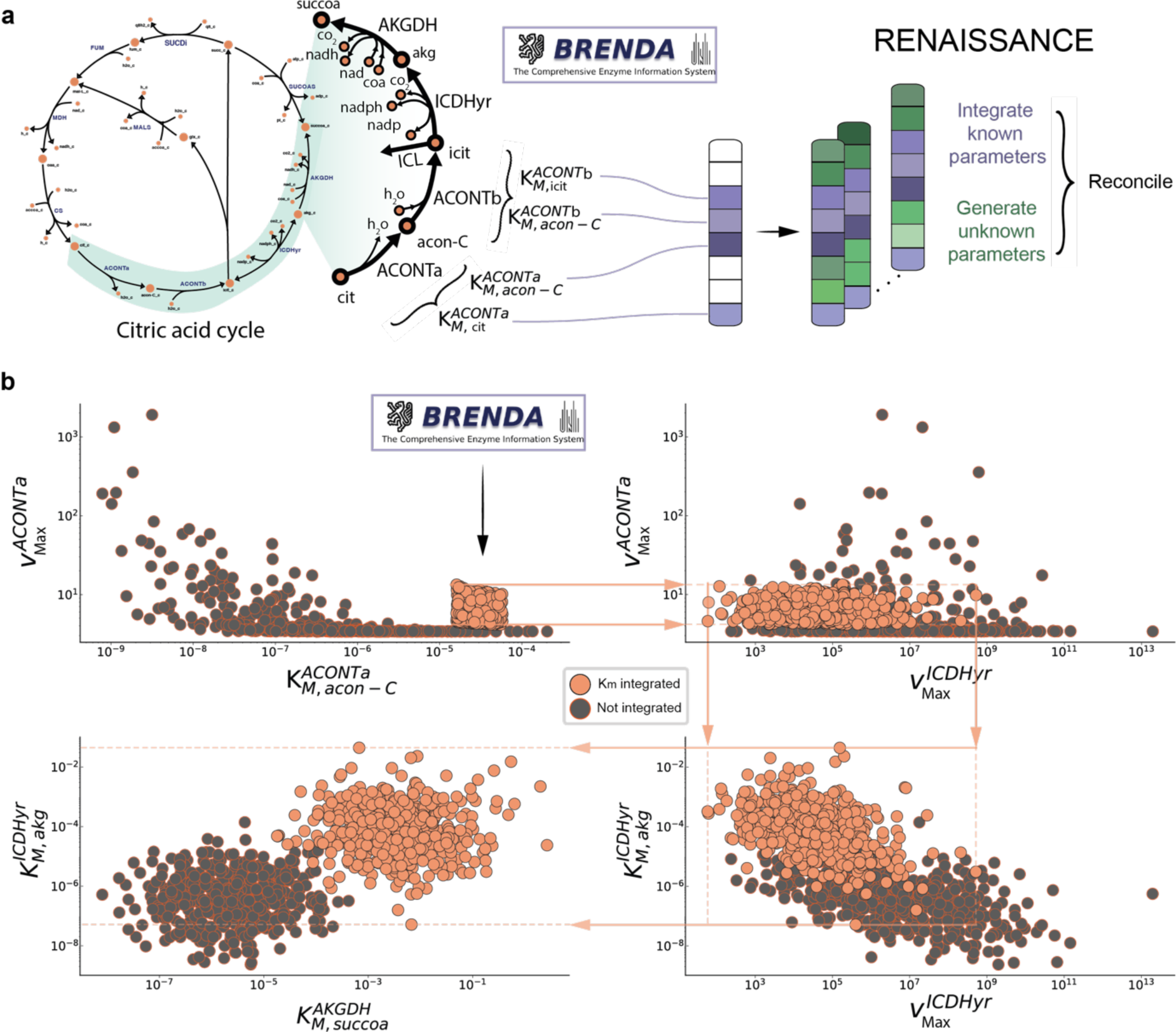
Integrated experimental *K*_*M*_ values for aconitase affect parameters of neighboring reactions. **a**, RENAISSANCE allows direct integration of *K*_*M*_ values from literature and reconciling them with unknown parameters that collectively lead to valid kinetic models. **b**, Propagation of the integrated *K*_*M*_ experimental data around aconitase (ACONTa, b) through the metabolic network. Comparison of RENAISSANCE generated values for: 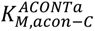 vs 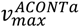 (upper left), 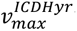 vs 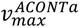 (upper right), 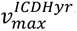 vs 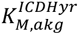 (lower right), and 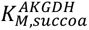 vs 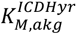 (lower left) when (i) no kinetic parameters (grey circles) (ii) 4 parameters are integrated (orange circles, Methods). Abbreviations: *ACONTa, b*: Aconitase, *ICDHyr*: Isocitrate dehydrogenase, *AKGDH*: 2-Oxoglutarate dehydrogenase, *acon-C*: Cis-Aconitate, *akg*: 2-Oxoglutarate, *succoa*: Succinyl Coenzyme-A.

Integration of *K*_*M*_ values of aconitase at a reaction level restricted the estimates of 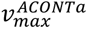 (Figure 4b, upper left). Due to the correlation in the *v*_*max*_ values throughout the network, restricting 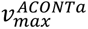 estimates through *K*_*M*_ integration constrained the estimated ranges of other maximal velocities, such as 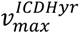 (Figure 4b, upper right). This restriction further affected downstream *K*_*M*_ values in the network, such as 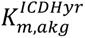 and 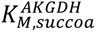 (Figure 4b, bottom left and right). These results suggest that integrating only a small amount of experimental data, localized to one enzyme (ACONTa, b), propagates throughout the metabolic network and alters the rest of the kinetic parameters.

We next enquired if RENAISSANCE improves its *K*_*M*_ estimates as the number of integrated experimental *K*_*M*_ values increases. We also examined how the localization of integrated *K*_*M*_ values, such as the integration of *K*_*M*_ values from the citric acid cycle, affects the estimation of *K*_*M*_ values in other subsystems of the metabolic network. Specifically, we integrated 10 random combinations of half (9) of the 17 available experimentally measured *K*_*M*_ values associated with the citric acid cycle (TCA) of *E. coli*, one combination at a time. For each of the 10 combinations, we obtained generators with a high incidence of valid models (> 90%) and generated 2000 of these models. In total, we generated 20,000 models containing 10 distinct combinations of the remaining 8 Michaelis constants to be estimated. This process ensured that each of the 17 Michaelis constants was integrated at least once and estimated at least once within the 10 combinations.

The comparison between the experimentally observed and the RENAISSANCE estimated range of TCA *K*_*M*_ values, quantified through the overlap (OS) between these two ranges (Fig 5d), showed that integrating *K*_*M*_s improves the estimates of the non-integrated individual *K*_*M*_s within the same subsystem (Fig 5a left, red bars, Supplementary Figure 9), compared to when no *K*_*M*_ values are integrated (black diamonds, Supplementary Figure 15). Indeed, significant improvement was observed in the predictions of 16 out of the 17 *K*_*M*_ values in TCA when experimental values of *K*_*M*_ were integrated. The average prediction accuracy for the entire subsystem also increased (Fig 5a right, red bars) compared to the case with no integration of experimental *K*_*M*_s (blue bars). A similar analysis was conducted for other subsystems, Pentose Phosphate Pathway (PPP), Glycolysis (gly), Anaplerotic reactions (anpl), Shikimate pathway (shkk), and Pyruvate metabolism (pyr), and consistently, estimates of *K*_*M*_ values within the same subsystem improved upon the integration of experimental information for all the cases (Fig 5a, right, Supplementary Figure 10). These findings indicate that integrating experimental information may improve prediction accuracy beyond the subsystem level.

**Fig. 5.**
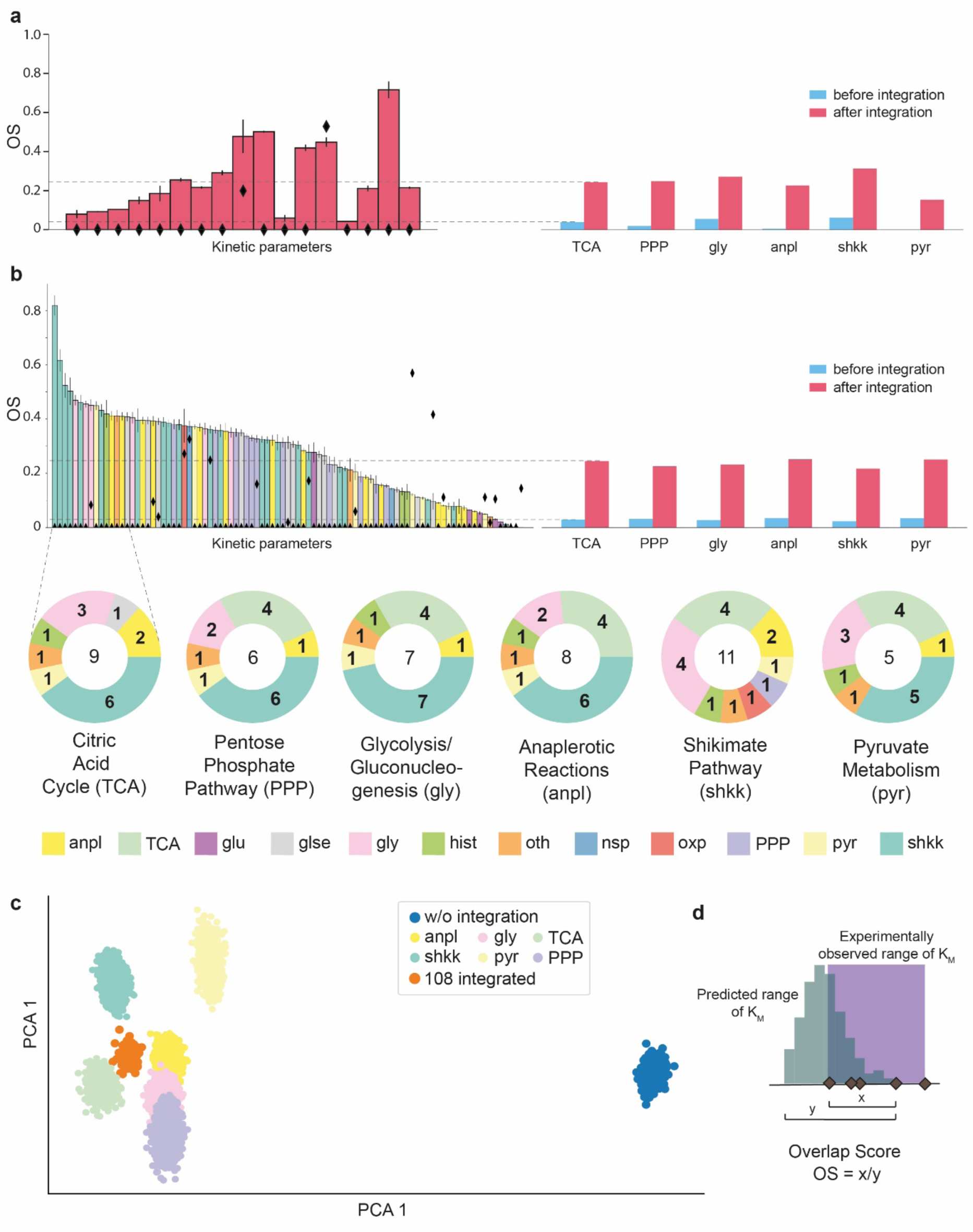
Integrating experimental kinetic information improves other parameter estimates: **a**, Left: The overlap scores (OS, defined in d) of RENAISSANCE estimates for 17 experimentally available Michaelis constants, *K*_*M*_s, from the citric acid cycle (red). The estimation process involved integrating experimental values for 9 out of the 17 parameters and estimating other model parameters including the remaining 8 parameters from TCA using RENAISSANCE. This procedure was repeated 10 times with a different random combination of 9 parameters. The black diamonds represent the OS of the estimates without integrating experimental data. The error bars indicate the standard error in the OS over 10 combinations of integrated parameters. Right: The mean OS for the estimated *K*_*M*_s within metabolic subsystems before integration (blue) and after integration (red) of experimentally measured *K*_*M*_s. The estimation process for each subsystem was conducted in a similar manner as in a. **b**, Upper Left: The overlap scores (OS) of RENAISSANCE estimates for 91 experimentally available *K*_*M*_s not belonging to TCA when 10 different combinations of 9 (out of 17) *K*_*M*_s from TCA were integrated. The black diamonds represent the OS without integration. The error bars represent the standard error in OS. Upper Right: The mean OS of the estimated *K*_*M*_s from all metabolic subsystems except for the subsystem labeled on the x-axis before integration (blue) and after integration (red) of *K*_*M*_s from the labeled subsystem. Bottom: The metabolic subsystems of the top 15 *K*_*M*_s with the highest increase in OS when Michaelis constants from the labeled subsystem were integrated. The numbers within the donut plots indicate the number of integrated experimental *K*_*M*_ values from the labeled subsystem. **c**, The 2-component PCA^48^ representation of the 276 (out of 384) experimentally unverified kinetic parameters with and without integrating experimentally measured Michaelis constants. **d**, The Overlap Score (OS) of an estimated *K*_*M*_ is calculated as the ratio between (x), the overlap of the RENAISSANCE predicted (teal) and the experimentally observed range (violet), and (y), the total range of the RENAISSANCE prediction. *Abbreviations: anpl* **–** Anaplerotic reactions, *TCA* – Citric Acid Cycle, *glu* – Glutamate Metabolism, *gly* – Glycolysis/Gluconeogenesis, *glse* – Glycine & Serine metabolism, *hist* – Histidine Metabolism, *nsp* – Nucleotide Salvage Pathway, *oxp* – Oxidative Phosphorylation, *PPP* – Pentose Phosphate Pathway, *pyr* – Pyruvate metabolism*, shkk* – Shikimate pathway, oth – Other reactions.

Inspecting the distributions of the generated *K*_*M*_s that were not part of the TCA subsystem revealed that the predictions for a vast majority of these *K*_*M*_s (85 out 91) improved upon the integration of TCA *K*_*M*_s (Fig 5b, upper left, colored bars) compared to the case where no *K*_*M*_s were integrated (black diamonds). Similarly, the mean overlap score (OS) of the entire set increased (Fig 5b, upper right). We then examined the top 15 *K*_*M*_s that exhibited the most significant improvement in their estimates and determined the metabolic subsystem in which they are located. The integration of experimental *K*_*M*_ values from TCA yielded the most significant improvement in the estimates of the shikimate pathway (6 in the top 15), followed by glycolysis (3 out of 15) and anaplerotic reactions (2 out of 15) (Fig 5b, leftmost donut plot). Interestingly, a similar analysis conducted by integrating *K*_*M*_s from other subsystems showed that the estimates from these three subsystems (shkk, gly, and anpl) consistently yielded the most significant improvement (Fig 5b, Supplementary Figure 11). These results provide evidence that RENAISSANCE effectively incorporates experimental kinetic data from a specific subsystem of the metabolic network, resulting in improved parameter estimates across the entire network.

We further examined the impact of integrating experimental kinetic data on parameters that lack verifiable experimental measurements, which accounted for 276 out of 384 *K*_*M*_s. To obtain a qualitative assessment of the effects of integration, we employed PCA^45^ to visualize the RENAISSANCE predictions for these unknown *K*_*M*_s (Figure 5c). The analysis revealed notable shifts in the estimates of these *K*_*M*_s when experimental data was integrated compared to the case where no data was integrated (Fig 5c, blue cluster). Additionally, the estimates for the cases with integrated experimental data exhibited greater similarity than those without integration. We provide the generator, trained to incorporate all 108 *K*_*M*_s available from BRENDA (Supporting Figure 12), in the Supporting data.

These results suggest that integrating experimental kinetic information reduces quantitative uncertainties in the intracellular metabolic state of the cell, allowing RENAISSANCE to make more informed predictions on the dynamic properties of the entire metabolic network. We anticipate that the inclusion of new experimental data and its subsequent integration will enhance the predictive capabilities of RENAISSANCE even further.

## Discussion

Metabolism plays a defining role in shaping the overall health of living organisms. A reprogrammed or altered metabolism is not only associated with the most common causes of death in humans – cancer, stroke, diabetes, heart disease, and others – but is also related to many congenital diseases^49^. Thus, a better understanding of metabolic processes is crucial to accelerate the development of new drugs, personalized therapies, and nutrition. Biotechnological advances like the bioproduction of industrially essential compounds and environmental bioremediation also hinge on our ability to describe cellular metabolism accurately.

Kinetic models provide the most thorough mathematical representation of metabolism. The efficient construction of these models will open new possibilities for various biomedical and biotechnological applications. However, acquiring the parameters of these models with traditional kinetic modeling approaches is computationally expensive and arduous^15,34^. Several machine learning methods were recently proposed for more efficient kinetic model generation, including iSCHRUNK^32,33,50^ and REKINDLE^34^. REKINDLE, in particular, has demonstrated remarkable gains in model generation efficiency by using generative adversarial networks (GANs)^51^. Nevertheless, existing kinetic modeling approaches were required to create the data needed for the GAN training. The proposed RENAISSANCE framework retains the model generation efficiency of REKINDLE without the need for training data.

In its conception, RENAISSANCE can parameterize kinetic models to satisfy a broad range of biochemical properties or physiological conditions. For example, it can parameterize models reproducing experimentally observed fermentation curves or drug adsorption patterns. Herein, we use RENAISSANCE to parameterize kinetic models to be consistent with an experimentally observed steady state. This approach to model construction was introduced within the ORACLE conceptual framework^52–55,50^, which parameterizes kinetic models by unbiased sampling. In contrast, in RENAISSANCE, we leverage machine learning to perform stratified sampling biased toward kinetic models producing metabolic responses over time with timescales^37^ matching experimental observations in studied organisms. Due to its capability to bias parameter sampling toward desired model properties, the proposed framework substantially improves model construction efficiency, enabling comprehensive studies of multiple physiological conditions.

RENAISSANCE can train model generators on a standard workstation in 3 to 20 minutes (Supplementary Note 5). Once trained, the generators generate ∼1 million models in 15-20 seconds, making this framework several orders of magnitude more efficient than traditional sampling-based kinetic approaches. RENAISSANCE also does not require specialized hardware to execute. The proof-of-concept applications shown here demonstrate RENAISSANCE’s applicability to a broad range of studies. In this work, we deployed RENAISSANCE to parameterize valid models of metabolism consistent with an experimentally observed steady-state, with validity being characterized by the biological relevance of their timescales. However, conceptually, any other requirement can be imposed or data used, such as consistency with knockout studies or time series from drug absorption trials.

As RENAISSANCE is agnostic to the nature, range, and number of the parameters it needs to generate, it is straightforward to adjust the framework to meet the specific demands the models need to satisfy. The parameters this framework can handle are not restricted to Michaelis constants only and can include other kinetic parameters, such as enzyme saturations^52^ and enzyme states^56^, and other unknown quantities in the studied system, such as metabolite concentrations.

Crucially, given proteomic data, RENAISSANCE can predict unknown enzyme turnover number, *k*_*cat*_, values and consolidate them with experimentally measured *k*_*cat*_ values from databases such as BRENDA and SABIO-RK^57^. As such, it represents a valuable complement to current machine learning methods that estimate *k*_*cat*_ values directly^58–60^.

In summary, we provide a fast and efficient framework that leverages machine learning to generate biologically relevant kinetic models. The open-access code of RENAISSANCE will facilitate experimentalists and modelers to apply this framework to their metabolic system of choice and integrate a broad range of available data.

## Methods

### *E. coli* model structure and data integration

The kinetic model structure is based on a previous study by Narayanan et al^38^. The reduced stoichiometry was obtained using redGEM^61^ and lumpGEM^62^, and it includes core carbon pathways such as glycolysis, the pentose phosphate pathway (PPP), the tricarboxylic cycle (TCA), anaplerotic reactions, the shikimate pathway, glutamine synthesis, and a lumped reaction for growth. The anthranilate phosphoribosyltransferase (ANPRT) was removed to tailor the general *E. coli* model to strain W3110 *trpD9923*. The resulting model structure had 113 mass balances, including one for biomass accumulation, involving 123 reactions parametrized with 507 kinetic parameters including 384 *K*_*M*_*s* and 123 *v*_*max*_*s* (Supplementary Figure 4). Further details on the kinetic model structure can be found in Supplementary Data 1 of Narayanan et al^38^.

A context-specific model of W3110 *trpD9923* was created by integrating metabolomics and fluxomics data from previous experimental studies. The lower bounds of the growth rate (0.26 hr^−1^) and anthranilate secretion rate (0.14 mmol/gDW/hr) were set to the reported values from Balderas-Hernandez et al^39^, and the glucose uptake rate was adjusted to be consistent with the secretion and growth rates. Extracellular metabolites neither found in the media nor listed as secreted were assigned upper bounds on their secretion rates of 0.01 µM gDW^−1^hr^−1^ and concentrations of 1μM, whenever possible. Intracellular metabolite concentrations reported in Park et al.^44^ were constrained to be within 2-fold of the reported values. Next, constraints were imposed on thermodynamic variables, calculated using the Group Contribution Method^42,43^, to ensure that the sampled flux directionalities and metabolite concentrations were consistent with the second law of thermodynamics.

5000 sets of steady-state profiles consistent with the integrated data were then sampled from this context-specific model using thermodynamics-based flux balance analysis implemented in the pyTFA tool^40^. Each steady-state profile comprises metabolite concentrations, metabolic fluxes, and thermodynamic variables. Once these profiles are available, we can generate kinetic models around each of these steady-states^53,55,27^ using the RENAISSANCE framework. We used the profile with index 1712 as input for RENAISSANCE in all studies, except for the study detailed in Section ‘Characterizing the intracellular states of E. coli metabolism’, where all 5000 steady states were employed (Figure 3).

At present, RENAISSANCE allows for integrating transcriptomics and proteomics data at the steady-state level. When computing the steady-state profiles around which the kinetic models are built, REMI^63^ and TEX-FBA ^64^ tools allow simultaneous integration of transcriptomics, metabolomics, and fluxomics data. Other tools for integrating transcriptomics are provided elsewhere^65^. As discussed in Sanchez & Zhang et al.^9^, proteomics data can be integrated by imposing the upper bounds on the fluxes through the expression 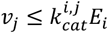, where *k*_*cat*_ is the turnover number, and *E* denotes enzyme concentration. The transcriptomics and proteomics data were not available for the present study.

### Determining the validity of kinetic models

The Jacobian matrix and its eigenvalues are used in the control^66^ and nonlinear dynamics theory^67^ to analyze the stability and behavior of a nonlinear dynamical system in the vicinity of an equilibrium point by performing linearization. The Jacobian matrix is derived by taking the partial derivatives of a set of differential equations with respect to the state variables. The sign of the Jacobian eigenvalues provides information on the local stability of the generated models, where a model is locally stable if the real parts of all eigenvalues are negative^52^. The inverse of the real part of the largest eigenvalue of the Jacobian defines the dominant time constant of the linearized system.

The time constant defines the time required for the system response to decay to 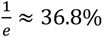 of its initial value. The dominant time constants allow us to characterize the model dynamics - small time constants characterize fast metabolic processes such as glycolysis and electron transport chain. In contrast, polymerization processes involving the synthesis of DNA, RNA, and proteins typically occur at slower time scales.

In this context, we consider a kinetic model valid (biologically relevant) if all time constants of the model response are consistent with the experimental observations of the studied organism.

To ensure that a perturbation of the metabolic processes settles within 1% of the steady state before cell division, the dominant time constants of the model response should be five times faster than the cell’s doubling time^38^. The biochemical response should also have a characteristic time slower than the timescale of proton diffusion within the cell^34^. With these properties, models can reliably describe the experimentally measured metabolic responses.

The doubling time of the *E. coli* strain used in this study is *t*_*doubl*@*n*9_ = 134 *mins*, which corresponds to a growth rate of 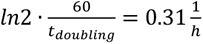. Therefore, the dominant time constant of the model’s responses should be smaller than one-fifth of the doubling time (26.8 *mins*). Here, we imposed a stricter dominant time constant of 24 minutes, corresponding to an upper limit of *Re*(*λ_i_*) < −2.5 (or −60/24), on the real parts of the eigenvalues, *λ_i_*, of the Jacobian. All kinetic parameter sets resulting in the model obeying this constraint are labeled valid and the rest invalid.

### Assigning rewards to determine fitness in RENAISSANCE

In RENAISSANCE, we employ deep neural networks known as generators to produce kinetic parameter sets for a given metabolic model structure. Technically and structurally, these neural networks are similar to those in generative adversarial networks (GANs)^51^ or other deep generative algorithms such as variational autoencoders (VAEs)^68^. The key distinction lies in the training methodology for this neural network. Generative adversarial networks (GANs) or variational autoencoders (VAEs) rely on explicit training data. For instance, in a prior study^34^, we used kinetic parameter sets derived from traditional kinetic modeling methods for training. Unlike traditional gradient-based deep learning methods, which rely on training data to train a neural network, RENAISSANCE employs the Natural Evolution Strategy, NES (Supplementary Note 1), which only requires a scoring function.

To optimize the weights of the generator network, the NES algorithm produces a population of candidate solutions to an optimization problem and assigns a fitness score to each candidate solution (Figure 1a). The algorithm uses the fitness scores of the current solutions to generate the next generation of candidate solutions, which are likely to have better fitness scores than the current generation. The iterative procedure stops as soon as the obtained solutions are satisfactory. This method is particularly advantageous in scenarios where the fitness landscape is complex, non-differentiable, or unknown, and it avoids the need for backpropagation or direct gradient computation.

The NES algorithm includes several steps:

Step I: NES collects the rewards of all the generators (neural networks) in a generation. In our case, the reward is the percentage of relevant generated models (thus, the reward for each neural network or generator in a generation is between 0 and 1).

Step II: The rewards for each generation member are normalized by subtracting the mean and dividing by the standard deviation of all the rewards in the generation. The normalization ensures that the update direction depends on how each member’s performance compares to the average rather than on absolute reward values. Step III: For each weight in the generator neural network, the algorithm computes an update proportional to the dot product between the population’s perturbations (the added noise for generating new individuals) and the normalized rewards. If a perturbation consistently results in higher rewards, it will exert a more significant influence on the direction of the weight update. Thus, the reward determines the selection of the ‘best’ generator. Step IV: The update to each weight is scaled by the learning rate and inversely scaled by the population size and the noise level (σ). Scaling prevents drastic changes that might destabilize the learning process (Supplementary Note 2).

The design objective of the conducted studies is to maximize the occurrence of biologically relevant kinetic models. In case all generators of the current generation fail to generate valid models, the generator producing models closest to the cut-off eigenvalue (−2.5) is rewarded higher, thus having a more significant impact on the weight of the parent generator in the next generation. The implementation of this concept is as follows.

To calculate the local gradient estimate, NES requires an objective function, *F*, to evaluate the fitness of each generator network, *G*. In our study, we use the incidence of the generator, *I*(*G*), as the objective function, which is defined as the fraction of the generated models that are relevant (0 ≤ *I*(*G*) ≤ 1). Thus, generator networks with a higher incidence of relevant models are ‘fitter’ than those with low incidence and have a higher weight in determining the parameters of the seed generator network for the next generation. In many cases, we observed that initially the generator neural networks do not generate any relevant models, and thus the optimization does not proceed as the fitness is always 0. To mitigate this, we added a sigmoidal term defined as follows,

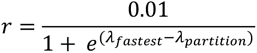

where, *λ_f_*_*astest*_ corresponds to the smallest maximal eigenvalue of the generated models and *λ*_*partition*_ is the maximal eigenvalue partition that determines the relevancy of the kinetic model. In this study, *λ*_*partition*_ = −2.5 (see previous section). This term rewards generators that generate models with dynamics closer to the relevant range more than those that generate models with slower, irrelevant, or unstable dynamics. This effectively pushes the optimization process toward finding generators that generate relevant models. So, the overall reward, *R*, for a generator, *G*, can be summarized as

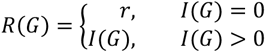

For the large-scale analysis of intracellular states (Fig. 3), the fitness for NES was no longer the incidence of the generators but the fastest possible dynamic for the models generated by a given generator. Thus, the reward was changed suitably as follows,

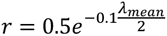

where *λ*_*mean*_ is the mean of the 10 fastest maximum eigenvalues (Supplementary figure 6) generated by a generator (out of 100 for this case study). This reward function ensured that the generators which generated models with more negative maximum eigenvalues (faster linearized dynamics, λ_NOP_) were rewarded more than the others.

### Hyperparameter tuning of RENAISSANCE

The hyperparameters of the NES algorithm used in RENAISSANCE are (i) the population size, *n*, determining the number of generator networks initiated/created and evaluated in each RENAISSANCE generation; (ii) search radius, *σ*, representing the level of noise injected into the weights of the parent generator in each generation; (iii) learning rate, *⍺*, determining the step size taken by the optimizer in the gradient space, i.e., it determines the magnitude of the updates to the generator neural network weights during each iteration of the optimization process; and (iv) learning rate decay, *d*, representing the rate at which the learning rate decreases at each generation, helping the optimization process to converge more effectively. In this study, we tuned the hyperparameters of RENAISSANCE to maximize the incidence of valid models (Supplementary Notes 3 and 4).

The optimal set of hyperparameters found after grid search is as follows: the population size of the generator networks, *n* = 20, the noise level in generating the agent population from the mean optimal weights in each generation, *σ* = 10^−2^, the learning rate of the gradient step, *α* = 10^−3^, and the decay rate of learning, *d* = 5%. In addition, the generated *K*_*M*_s were constrained strictly between {1.3 × 10^−11^, 20} to accurately represent experimentally measured *K*_*M*_ values as curated in the BRENDA database^48^.

### Generator neural networks

The generator neural networks were composed of three layers with 1,076,352 parameters: layer 1, Dense, with 256 units, Dropout (0.5); layer 2, Dense, with 512 units, Dropout (0.5); and layer 3, Dense, with 1024 units, Dropout (0.5). All software programs were implemented in Python (v3.6). Neural networks were implemented using the TensorFlow library^69^ (v2.3.0).

### Dimension reduction and visualization of steady states

For generating Fig. 3 a, d, f (left), the following steps were followed: I) the steady state matrix (consisting of 1127 features) was subjected to principal component analysis (PCA)^45^. II) The components of PCA that contributed to over 99% of the total expected variance were reduced to 2 dimensions using t-SNE^46^. III) The t-SNE components

{*x*_*q*_, *x*_*p*_} were then subjected to polar coordinate transformation as follows,

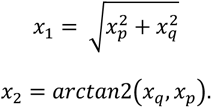

{*x*_1_, *x*_2_} were then plotted to generate the figures.

### Integrating known kinetic parameters in BRENDA

If there were multiple experimentally measured values for a single *K*_*M*_ in BRENDA, we took the geometric mean (*K*_*M*,*exp*_) of the different values and added an experimental error rate of ± 20% to *K*_*M*,*exp*_. The same error rate was applied if there was only one recorded experimental value, *K*_*M*,*exp*_. Then the value of an integrated *K*_*M*_ was sampled uniformly from the range *K*_*M*,*exp*_± 20% when integrated into RENAISSANCE for the training process and generation.

## Abbreviations

NES: Natural Evolution Strategies
ORACLE: Optimization and Risk Analysis of Complex Living Entities
GEM: GEnome-scale Model
RENAISSANCE: REconstruction of dyNAmIc models through Stratified Sampling using Artificial Neural networks and Concept of Evolution strategies
REKINDLE: REconstruction of KINetic models of metabolism using Deep LEarning
TFA: Thermodynamics-based Flux Balance Analysis
ODE: Ordinary Differential Equations

## Data availability

The data supporting this study’s findings are publicly available in the Zenodo repository (https://doi.org/10.5281/zenodo.7628650 and the links therein).

## Code availability

A Python implementation of the RENAISSANCE workflow is publicly available at https://github.com/EPFL-LCSB/renaissance and https://gitlab.com/EPFL-LCSB/renaissance. The ORACLE framework is implemented in the SKimPy (Symbolic Kinetic models in Python)^62^ toolbox, available at https://github.com/EPFL-LCSB/skimpy.

## Acknowledgments

This work was supported by funding from the Swiss National Science Foundation grant 200021_188623, the European Union’s Horizon 2020 research and innovation programme under grant agreement 814408, Swedish Research Council Vetenskapsradet grant 2016-06160, and the Ecole Polytechnique Fédérale de Lausanne (EPFL).

## Author contributions

S.C., M.M., and L.M. designed the overall method and approach. V.H. and L.M. supervised the research. S.C. and L.M. developed the RENAISSANCE method. S.C., B.N., and M.M. designed the code. S.C., B.N., and L.M. analyzed the data. S.C. and L.M. wrote the manuscript. All authors read and commented on the manuscript.

