## Supplementary material for "Generative machine learning produces kinetic models that accurately characterize intracellular metabolic states"

### Supplementary Note 1: Natural Evolution Strategies for Neural Networks

**Introduction**

One of the key goals of artificial intelligence is to develop agents that can accomplish predefined tasks in a complex and often uncertain environment with many unknown constraints. Machine learning, and more recently deep learning, has been shown to be adept at performing such complex tasks, especially real-world data classification and generation[1]. However, deep learning, like many other machine learning methods, depends on the availability of training data. Consequently, Reinforcement Learning[2] (RL) has emerged as a promising paradigm where neural network agents are trained to perform specific tasks in a contained environment using Markov Decision processes, thus eliminating the need for training data. In RL, policy functions (neural networks) are trained by injecting stochasticity in the actions they take and keeping a score of the rewards attained from the environment by these actions. The policy function parameters are then tweaked using backpropagation, iteratively making actions with higher rewards more frequent. RL has been successfully applied to solve a plethora of problems in various fields, including biology[3], computer vision[4], software development[5], and finance[6]. Alternative approaches to solving RL problems that do not require backpropagation have also been proposed, like Direct Policy Search[7], Neuro-Evolution[8], and Evolution Strategies[9]. RENAISSANCE utilizes Evolution strategies as its optimization strategy.

**Black Box Optimization using Evolution Strategies (ES)**

In a recent study[10] by OpenAI, ES was shown to be as competitive as RL at solving similar tasks with some additional benefits (Supplementary Note 2). The version of ES used in this study is from a class of natural evolution strategies[10](NES). NES uses heuristics inspired by natural evolution to achieve its optimization goal: a population of parameter vectors (that directly determine the objective fitness) is perturbed and their objective function value ('fitness') is evaluated. The highest scoring parameters are then recombined to form the next generation of parameter vectors. This is done iteratively over 'generations' until desirable objective fitness is achieved.

Let $F$ denote the objective function which takes in a parameter vector θ to determine fitness $F(\theta)$. Let $p_\psi(\theta)$ denote the distribution of the perturbed parameters in NES algorithm, where the distribution $p$ is itself parameterized by ψ. Then, the objective of NES algorithm is to find ψ that maximizes the average objective value, $\boldsymbol{E}_{\theta\sim p_\psi}F(\theta)$, using stochastic gradient ascent. The gradient estimator in ψ denoted as, $\nabla_\psi \boldsymbol{E}_{\theta\sim p_\psi}F(\theta)$, can be rewritten as,

$$\nabla_\psi \boldsymbol{E}_{\theta\sim p_\psi}F(\theta) = \boldsymbol{E}_{\theta\sim p_\psi}\{\nabla_\psi log p_\psi(\theta)\}$$

For the special case where $p_\psi$ is as an isotropic multivariate Gaussian with mean ψ and fixed covariance $\sigma^2 I$ (as in this study), one can write $\boldsymbol{E}_{\theta\sim p_\psi}F(\theta)$ directly in terms of the mean parameter vector $\theta$,

$$\boldsymbol{E}_{\theta\sim p_\psi}F(\theta) = \boldsymbol{E}_{\varepsilon\sim \mathbf{N}(\mathbf{0},\mathbf{I})}\, F(\theta + \sigma\varepsilon)$$

Where $\varepsilon$ is noise sampled from a standard multivariate Gaussian and $\psi = 0$. Then, the gradient estimator can be calculated as,

$$\nabla_\theta \boldsymbol{E}_{\varepsilon \sim \mathbf{N}(\mathbf{0},\mathbf{I})} F(\theta + \sigma\varepsilon) = \frac{1}{\sigma} \boldsymbol{E}_{\varepsilon \sim \mathbf{N}(\mathbf{0},\mathbf{I})} F(\theta + \sigma\varepsilon)\varepsilon$$

With the gradient step now defined, the resulting NES algorithm stochastically perturbs the candidate parameter vectors, evaluates the resulting parameters against the objective, combines the results to calculate the stochastic gradient estimate and updates the parameters. The pseudo-code can be written as follows,

1. **Inputs:** Learning rate **α**, noise standard deviation **σ**, number of agents: **n**, initial parameters, $\boldsymbol{\theta_0}$
2. **For** $t = 0,1,2,3, \ldots$ **do**
3. . . . . . **Sample** $\boldsymbol{\varepsilon_1, \varepsilon_2, \varepsilon_3, \ldots, \varepsilon_n \sim \mathrm{N}(0, \sigma^2 I_n)}$
4. . . . . . **Compute objective**: $F(\theta + \sigma\boldsymbol{\varepsilon_1})$ for $i = 0,1,2,3, \ldots n$
5. . . . . . **Update:** $\boldsymbol{\theta_{t+1} = \theta_t + \frac{\alpha}{n\sigma} \sum_{i=1}^{n} F_i \varepsilon_i}$
6. **End For**

### Objective function for RENAISSANCE:

RENAISSANCE uses NES algorithm to evolve generator neural networks that can consistently generate relevant kinetic parameter sets given some kinetic structure (in the form of a system of ODEs, $d\boldsymbol{x}/dt = \boldsymbol{S}.\boldsymbol{v}(\boldsymbol{x}, \boldsymbol{k}, E, t)$ where $\boldsymbol{S}$ is stoichiometry of the metabolic network), steady state vector of fluxes, $\boldsymbol{v}$, and concentrations, $\boldsymbol{x}$, and a mathematically defined design objective. In our previous workflow for reconstructing kinetic models using generative adversarial networks, REKINDLE[12], we showed that feed-forward, fully connected generator neural networks are appropriate function approximators for kinetic models of metabolic networks due to their high non-linearity.

Generators, $G(\boldsymbol{w})$, in REKINDLE take a noise vector, $\boldsymbol{z}$, sampled from a multivariate standard Gaussian distribution as input and generates kinetic parameter vectors given the model constraints as follows,

$$G(\boldsymbol{z} \mid \boldsymbol{w}, \boldsymbol{S}.\boldsymbol{v}, \{\boldsymbol{x}, \boldsymbol{v}\}) = \boldsymbol{k}$$

where $\boldsymbol{w}$ are the parameters of the neural network that are optimized during training and relevancy of the kinetic parameters is determined by the design objective. In this study, the design objective is to obtain kinetic parameters whose dynamics satisfy biophysical time scales like *E. coli* doubling time and intracellular molecular diffusion limits. If these conditions are met, the kinetic parameter is termed relevant. We define the incidence of a generator, $\boldsymbol{I}(\boldsymbol{w})$ as the fraction of relevant kinetic parameter vectors being generated. We set this incidence as our objective function, $\boldsymbol{F}(.)$, for the NES algorithm, which searches for $\boldsymbol{w}_{opt}$ that maximizes $\boldsymbol{I}(\boldsymbol{w})$.

### Supplementary Note 2: Comparison between RL and ES

Both RL and ES algorithms develop agents that solves complex tasks by interacting with unknown environments that sends feedback on how close it is to the objective. We opted for ES over RL in this study for the following reasons[10]:

- **No need for backpropagation:** As ES directly the perturbs the weights of the neural network it does not relies upon backpropagation to get gradient estimates. This makes the overall code shorter and faster to execute. This also eliminates the need for specialized hardware like GPUs that most deep learning libraries use for faster computation.
- **Highly parallelizable:** The agents in ES can be evaluated independently of each other as they do not need communicate except for the final fitness score. Thus, the process of initiating an agent (randomized neural network), making the forward pass on the agent, evaluating the agent output and the assignment of the score from the environment can be compartmentalized neatly within individual CPU cores/ threads. This leads to linear reductions in computation times depending on the number of agents initiated and the number of CPU cores used.
- **Easier to set up:** In line with the arguments mentioned above, ES requires a standard desktop to operate with fewer hardware/software requirements compared to RL.
- **More intuitive:** ES is easier to understand compared to RL, where policy and learning parameters are abstracted and difficult to understand in terms of the overall optimization process.

### Supplementary Note 3: RENAISSANCE hyperparameters

There are several hyperparameters that dictate the optimization in RENAISSANCE method (Supplementary Figure 1). They can be categorized as follows,

1. Hyperparameters of NES: number of agents, $n$, noise level in generating the agent population from the mean parameter vector, $\sigma$, learning rate of the gradient step, $\alpha$, and the decay rate of learning, $d$.
2. Hyperparameters of the dynamics of the metabolic network: the allowed ranges for the kinetic parameters and the desired dynamic timescales.

As explained in Supplementary Note 1, RENAISSANCE uses Evolution Strategies (ES) in the backend to achieve the desired modelling objectives (Supplementary Figure 1a). ES has certain hyperparameters that need to be tuned and fixed prior to optimization. Their effect on the optimization process is explained as below,

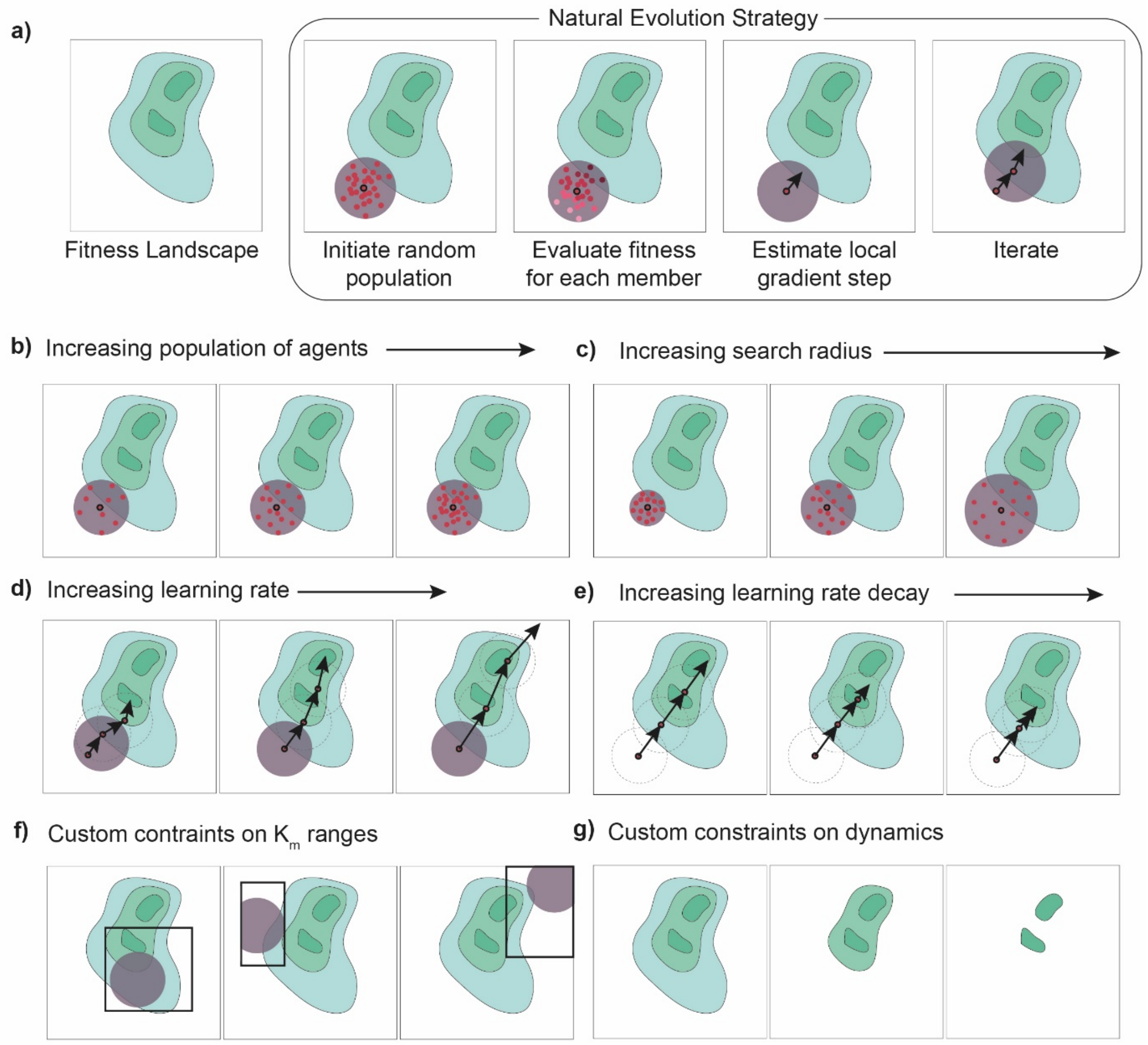


**Supplementary Figure. 1: Overview of the hyperparameters of RENAISSANCE:** a) Overview of Natural evolution strategies. Effect of increasing: a) Population size ($n$),. c) Search Radius ($\sigma$), d) Learning rate ($\alpha$), e) Learning decay rate ($d$) on the optimisation process. Effect of constraining f) Range of kinetic parameters, $K_m$ g) target range of dynamics on the fitness landscape.

1. Population size, $n$ : This hyperparameter determines the number of agents that will act on the environment (generate kinetic parameters) and be evaluated (checking the parameterized kinetic models for desired kinetic properties) in every iteration (Supplementary Fig. 1b). Having a large number of agents leads to a more thorough exploration of the landscape but also increases computational time.
2. Search radius, $\sigma$ : This hyperparameter determines the local radius from the mean agent in which the other agents will be sampled from, after the best fit parameter is updated using stochastic gradient ascent (Supplementary Figure 1c). Having a large search radius and small

population size will lead to an inadequate exploration of the landscape leading the optimization process to fail or collapse into a local optimum.

3. Learning rate, $\alpha$ : This hyperparameter determines the step size of the gradient step (Supplementary Fig. 1d). Having a too low learning rate will slow down the optimization process, while having a large learning rate might lead the optimization away from the global optimums, thus leading to a failure of the optimization process.
4. Learning rate decay, $d$ : This hyperparameter determines how fast the learning rate is decayed per iteration (Supplementary Fig. 1e). Having a fast decay will lead to the optimization slowing down too fast without exploring the landscape while having a very slow decay might make the optimization process skip away from the desired global optimum .

In addition to the hyperparameters inherent to NES, RENAISSANCE can be tuned for additional hyperparameters related to the dynamic properties of the model itself,

1. Ranges of $K_m$ vectors: The generated kinetic parameter vectors can be constrained to a specific range depending on the modelling objective and the system being studied. Constraining the ranges of the parameters leads to sections of the overall fitness landscape being accessible to RENAISSANCE, thus leading to different results (Supplementary Figure 1f).
2. Custom dynamic timescales: Depending on the modelling objective, different conditions can be imposed on the dynamics of the metabolic model, leading to a change in the optimization landscape and thus producing different results (Supplementary Figure 1g).

### Supplementary Note 4: RENAISSANCE hyperparameter tuning

The hyperparameters used in RENAISSANCE need to be tuned for good optimization results. As we did not find any reference for a tuned set of NES hyperparameters in literature, we tuned every hyperparameter individually by performing a grid search. We use the same metabolic models and the same objective function as the results in Fig. 2 (Methods). The hyperparameter tuning results are summarized in Supplementary Figure 2. We repeat every hyperparameter search 10 times with a randomly initialized population of agents (neural networks).

1. Learning rate ($\alpha$): We tested different learning rates ($10^{-1}, 10^{-2}, 10^{-3}, 10^{-4}$) while keeping the other hyperparameters fixed ($n = 20, \sigma = 10^{-2}, d = 5\%$). We observe that $\alpha = 10^{-3}$ performs significantly better than the other values (green line, Supplementary Fig. 2a). We thus choose our optimal learning rate, $\alpha_{opt} = 10^{-3}$.
2. Population size ($n$): We tested different population sizes of agents (10, 20, 50 and 100) while fixing the other hyperparameters ($\sigma = 10^{-2}, \alpha = 10^{-3}, d = 5\%$). We observe that the incidence of relevant models increases faster for higher $n$ (Supplementary Figure 2b). However, the computational time also increases with $n$ (Table 1). We choose $n = 20$ as our optimal population size, $n_{opt}$, as we get satisfactory results with reduced computational time.

3. Search Radius ($\sigma$): We tested different noise levels in the agent (generator neural network) parameters for creating the population for each iteration while fixing the other hyperparameters ($n = 20, \alpha = 10^{-3}, d = 5\%$). We observe that $\sigma$ significantly affects the incidence of relevant models (Supplementary Fig. 2c, left). We also plotted the final incidence after 25 generations for each $\sigma$ (Supplementary Fig. 2c, right). We observe that the incidence is maximum for $\log(\sigma) = -1.6\ to - 2$. Thus, we choose our optimal search radius, $\sigma_{opt} = 10^{-2}$.

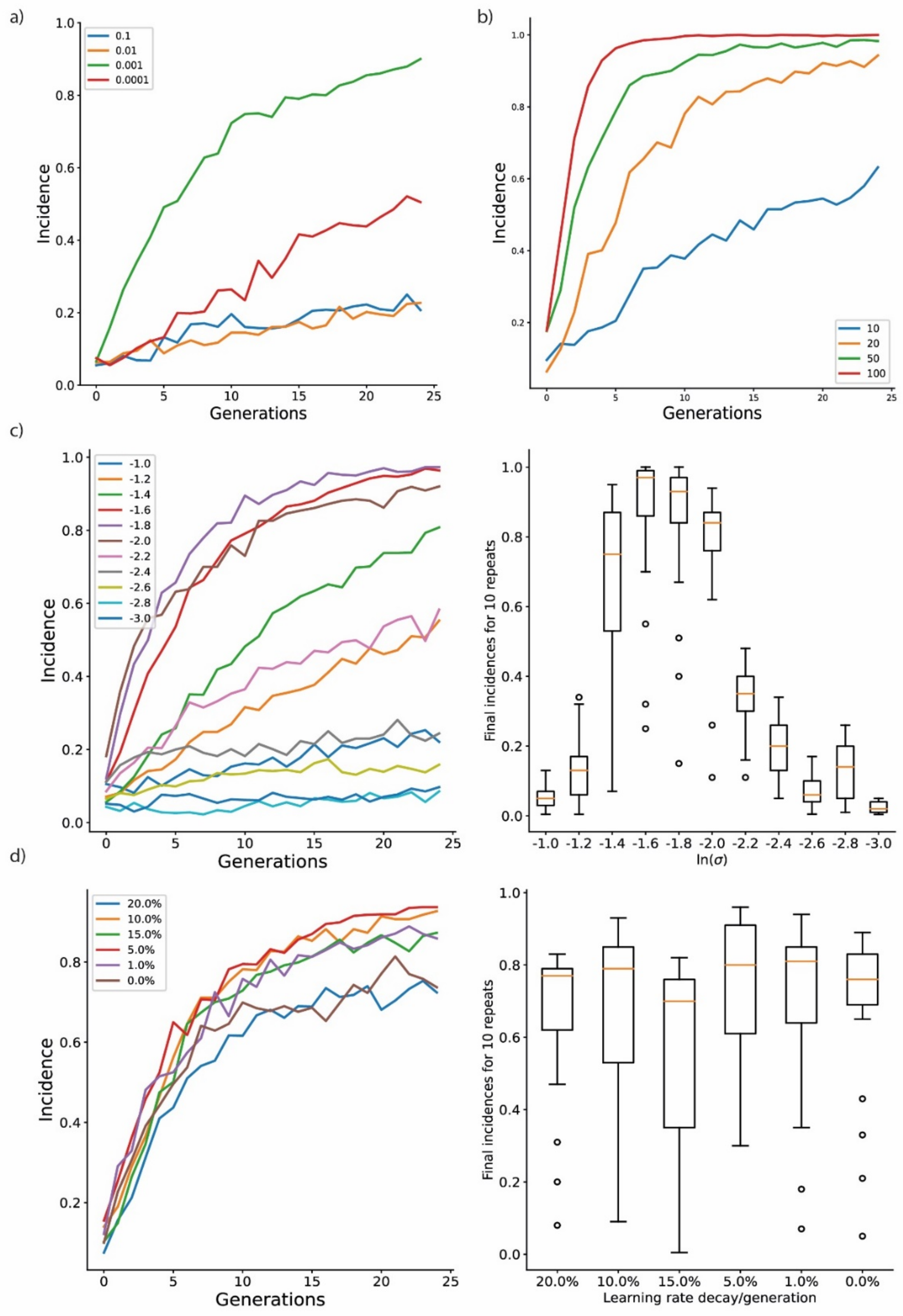

**Supplementary Figure. 2: Hyperparameter tuning of NES: a) Learning rate ($\alpha$):** Mean incidence of relevant models for different values of learning rate for 10 statistical repeats. **b) Population size ($n$) :** Mean incidence of relevant models for different values of population sizes for 10 statistical repeats. **c) Search Radius ($\sigma$): Left:** Mean incidence of relevant models for different values of search radii for 10 statistical repeats**. Right:** Final incidences reached for different values of search radii for 10 statistical repeats**. d) Learning decay rate ($d$): Left:** Mean incidence of relevant models for different values of learning rate decay per generation, for 10 statistical repeats**. Right:** Final incidences reached for different values learning rate decay per generation for 10 statistical repeats.

4. Learning decay rate ($d$): Rate We tested different decay rates for $\alpha$ ($0\%,\ 1\%,\ 5\%,\ 10\%,\ 15\%,\ 20\%$) while keeping the other hyperparameters constant ($n = 20, \sigma = 10^{-2}, \alpha = 10^{-3}$). We observed that having a small decay rate of $5\%$ outperforms having no decay at all (red line, Supplementary Fig. 2d). Having a higher decay rates does not lead to increase in performance. Thus, we choose our optimal decay rate, $d_{opt} = 5\%$.

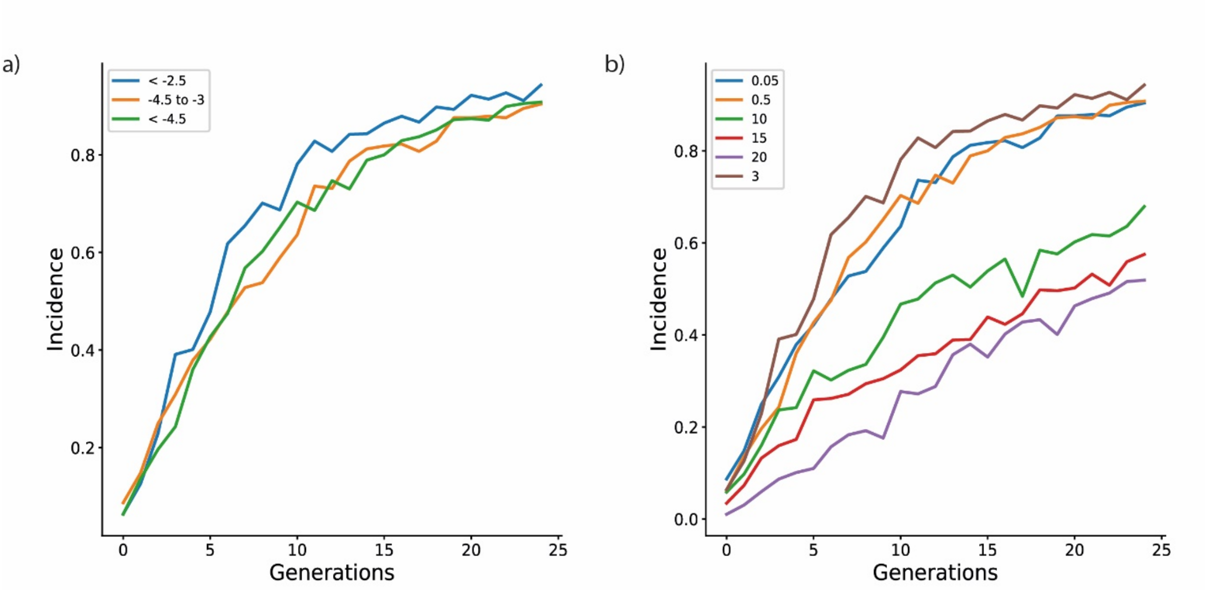


**Supplementary Figure. 3: a) Targeting different range of dynamics for same steady state:** Mean incidence of relevant models for different target range of eigenvalues for 10 statistical repeats. **b) Range of kinetic parameters, $K_m$:** Mean incidence of relevant models for different ranges of $K_m$ for 10 statistical repeats.

Next, we tested the effect of different constraints on the kinetic properties of the metabolic model on the optimization process of RENAISSANCE,

1. **Target different ranges of dynamics:** For the bulk of this study, a kinetic model is considered to be relevant if all the eigenvalues of the Jacobian are less than $-2.5$. However, this value can also be changed in RENAISSANCE to target dynamics that are faster ($< -2.5$) or slower ($> -2.5$). We tested RENAISSANCE by optimizing it for targeting different ranges of eigenvalues (less than $-2.5$, between $-4.5\ and\ -2.5$, less than $-4.5$) and hence corresponding dynamics. The results are summarized in Sup. Fig 3a. We observe that

optimization proceeds at the same rate regardless of target dynamic range of the kinetic models and RENAISSANCE successfully generates models that are consistently relevant.

2. **Range of kinetic parameters, $\boldsymbol{K_m}$:** We observed in literature[13] that depending on the organism and the specific strain being studied, the characteristic $K_m$s can take different range of values. The lower limit on the $K_m$s was always well constrained around $e^{-25} = 1.34 * 10^{-11}$. However, the upper limit ($\boldsymbol{K_{m,max}}$) varied between different organisms. For *E. coli*, the upper limit was around $e^1 = 2.7$. So, we chose an upper limit of $e^3 = 20.08$, to account for experimental errors for *in vitro* measurements listed in literature. We also tested the performance of RENAISSANCE for different values of the upper limit ($\ln(\boldsymbol{K_{m,max}}) = \mathbf{0.05, 0.5, 3, 10, 15, 20}$) respectively. The results are summarized in Sup. Fig. 3b. We observe that the rate of convergence of optimization is affected by the upper limit (faster for smaller values of $\boldsymbol{K_{m,max}}$).

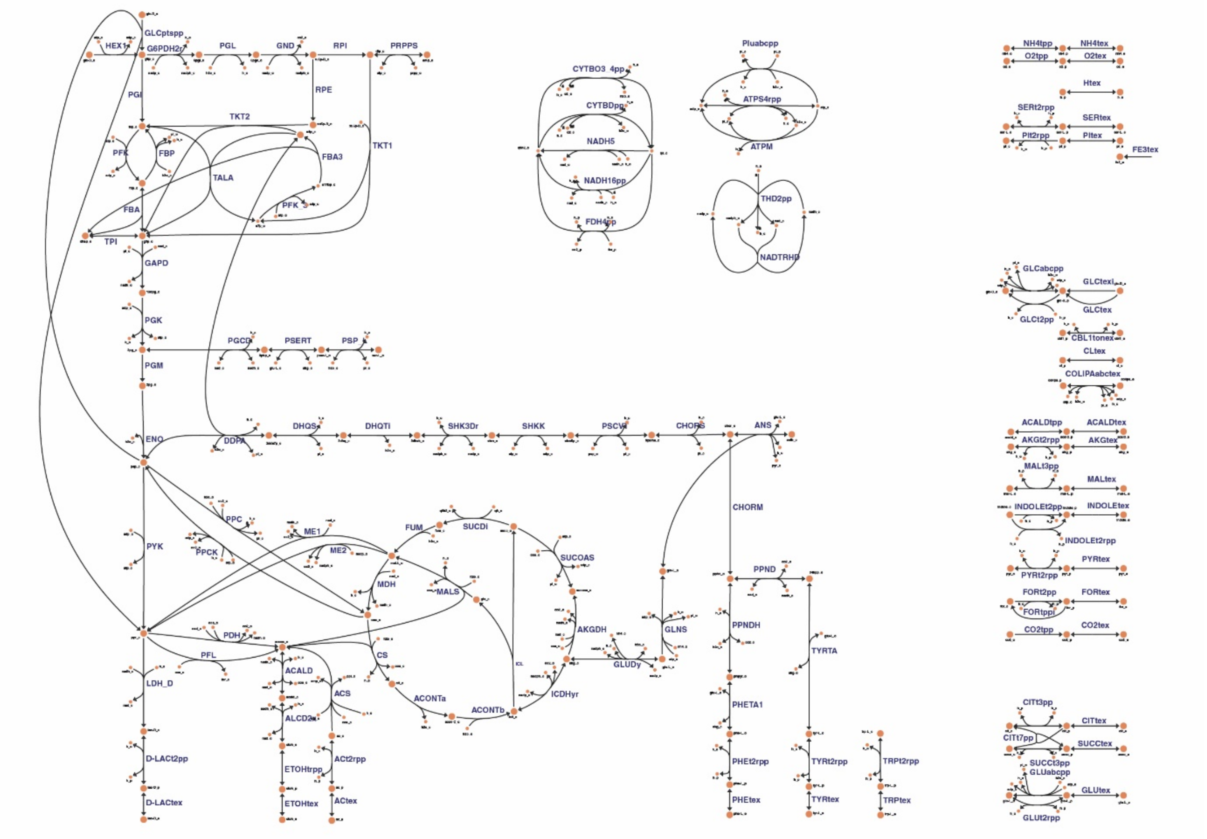


**Supplementary Figure. 4:** Network map of the *E.coli* metabolic model used for this study. The reactions are represented as black lines and the metabolites as orange circles. This map was generated using Escher web tool[14].

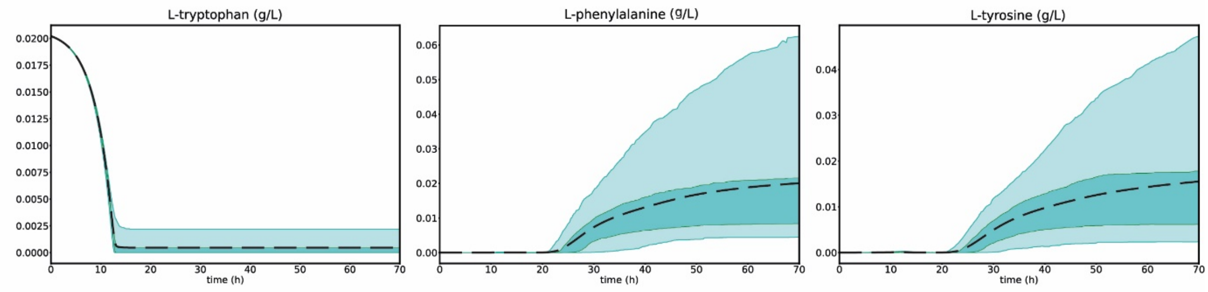


**Supplementary Figure. 5: (Left to Right)** The time evolution of (i) Tryptophan concentration (in g/L) (ii) Phenyalanine concentration (in g/l) (iii) Tyrosine concentration (in g/l) in the bioreactor simulation. The dashed black line represents the mean response, the dark cyan region corresponds to the 25$^{th}$-75$^{th}$ percentile and the light cyan region corresponds to 5$^{th}$-95$^{th}$ percentile of the ensemble of responses.

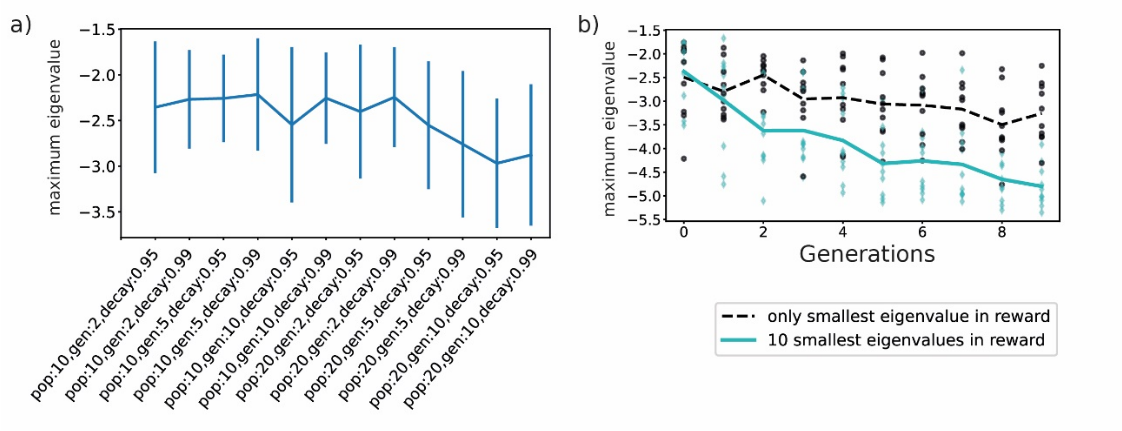


**Supplementary Figure. 6: (a)** Hyperparameter tuning of RENAISSANCE for finding the fastest dynamic (most negative maximum eigenvalue) for different values of Population size, $n$, number of generations, $g$, and learning rate decay, $d$ for 10 statistical repeats, for one random steady state out of 5000. $n = 20, g = 10$ and $d = 5\%$ gave the best results. **(b)** The maximum negative eigenvalue obtained by RENAISSANCE over 10 generations for 10 statistical repeats when considering (i) only the most negative eigenvalue of the generated models (black) (ii) average of the 10 most negative eigenvalues of the generated models (cyan) in the reward function (Methods).

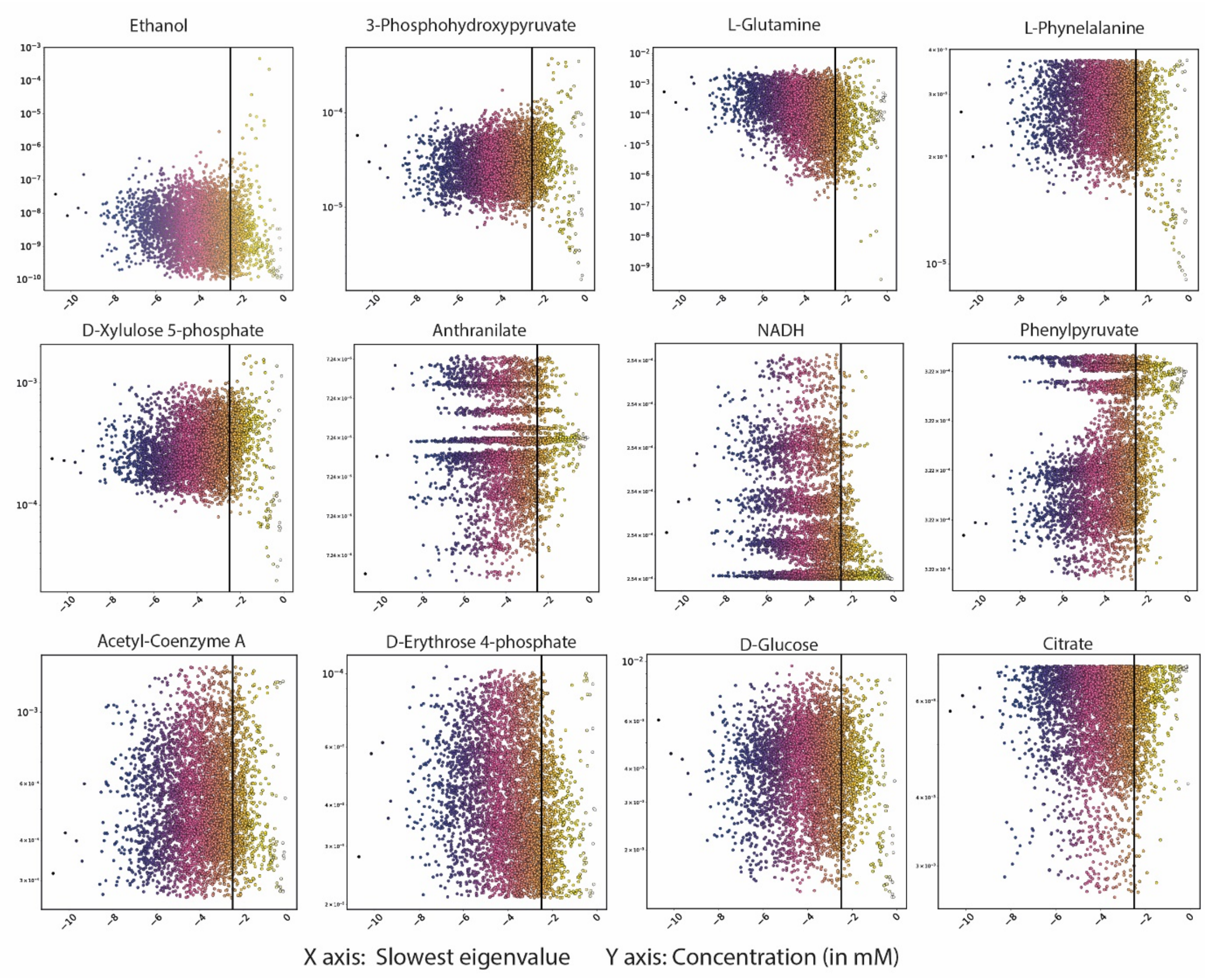


**Supplementary Figure. 7:** Concentration of metabolite (mM) in every steady state vs. the fastest linearized dynamic achieved by RENAISSANCE in every steady state for following metabolites, Top Panel: (left to right) Ethanol (etoh), 3-Phosphohydroxylpyruvate (3php), L-Glutamine (gln-L), L-Phenylalanine (phe-L) Middle Panel (left to right): D-Xylulose 5-phosphate (xu5p-D), Anthranilate (anth), NADH (nadh), Phenylpyruvate (phpyr). Bottom Panel (left to right): Acetyl-Coenzyme A (accoa), D-Erythrose 4-Phosphate (e4p), D-Glucose (glc-D), Citrate (cit). The black vertical line represents that cutoff for the linearized dynamic to relevant. We observe that for some of the metabolites, fast dynamics (blue/pink dots) are achieved only in certain continuous ranges of concentrations (etoh, 3php, gln-L, phe-L, xu5p). For some metabolites, fast dynamics are achieved in discrete ranges of concentrations (anth, nadh, phpyr). Some metabolites allow fast dynamics over the whole thermodynamically feasible range (accoa, e4p, glc-D, cit).

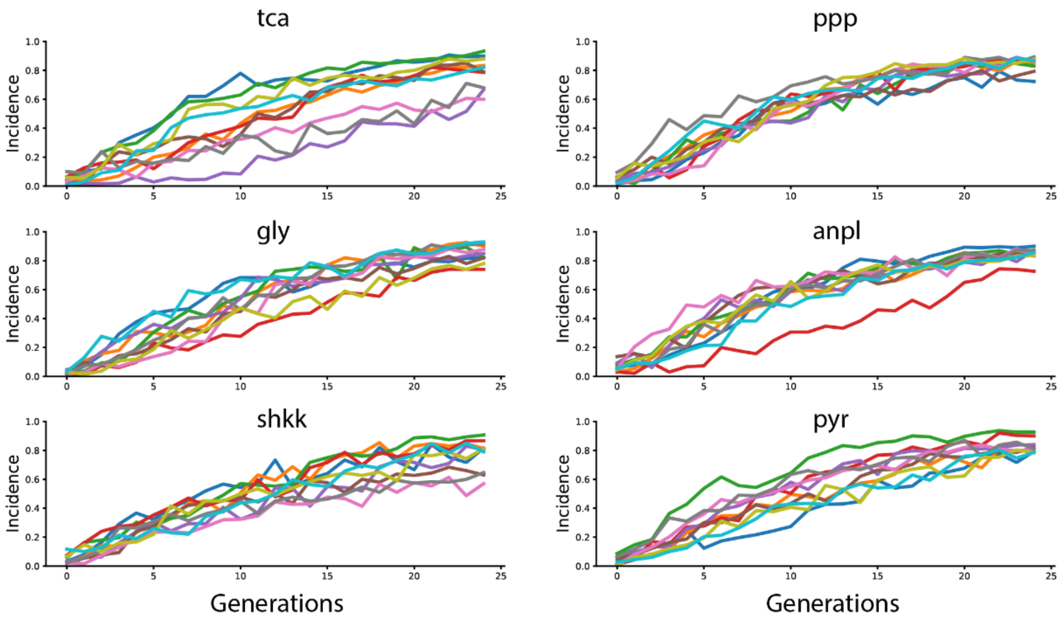


**Supplementary Figure. 8:** Mean incidence of relevant models over generations for 10 different random combinations (3 repeats for each combination) of $K_M$s from, Abbreviations: **tca:** Citric Acid Cycle, **ppp:** Pentose Phosphate Pathway, **gly:** Glycolysis/Gluconucleogenesis, **anpl:** Anaplerotic reactions, **shkk:** Shikimate pathway, **pyr:** Pyruvate metabolism.

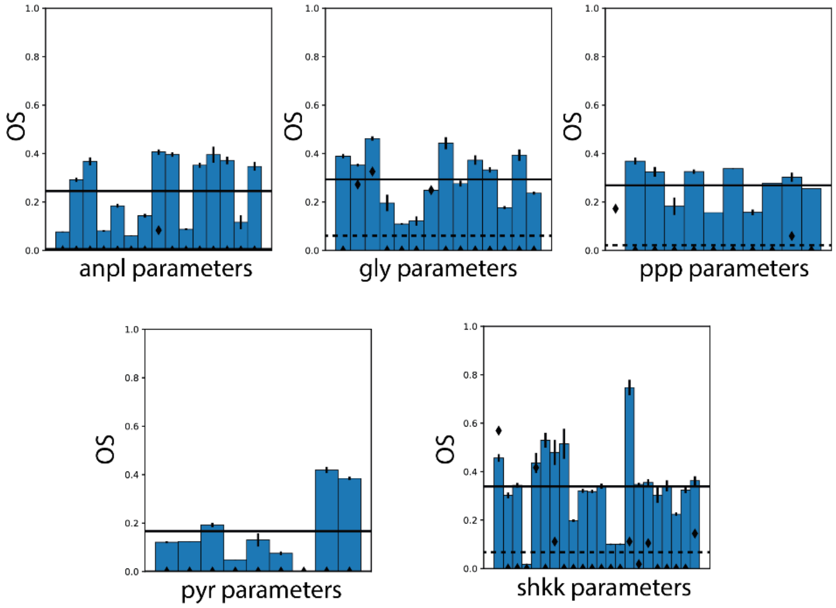


**Supplementary Figure. 9:** The mean overlap scores (OS) of $K_M$s belonging to a metabolic subsystem when $K_M$s from the same subsystem are integrated into RENAISSANCE (blue bars). The error bars represent the standard error in the OS. The black diamonds represent the OS when no $K_M$s are integrated. Abbreviations: **ppp:** Pentose Phosphate Pathway, **gly:** Glycolysis/Gluconucleogenesis, **anpl:** Anaplerotic reactions, **shkk:** Shikimate pathway, **pyr:** Pyruvate metabolism.

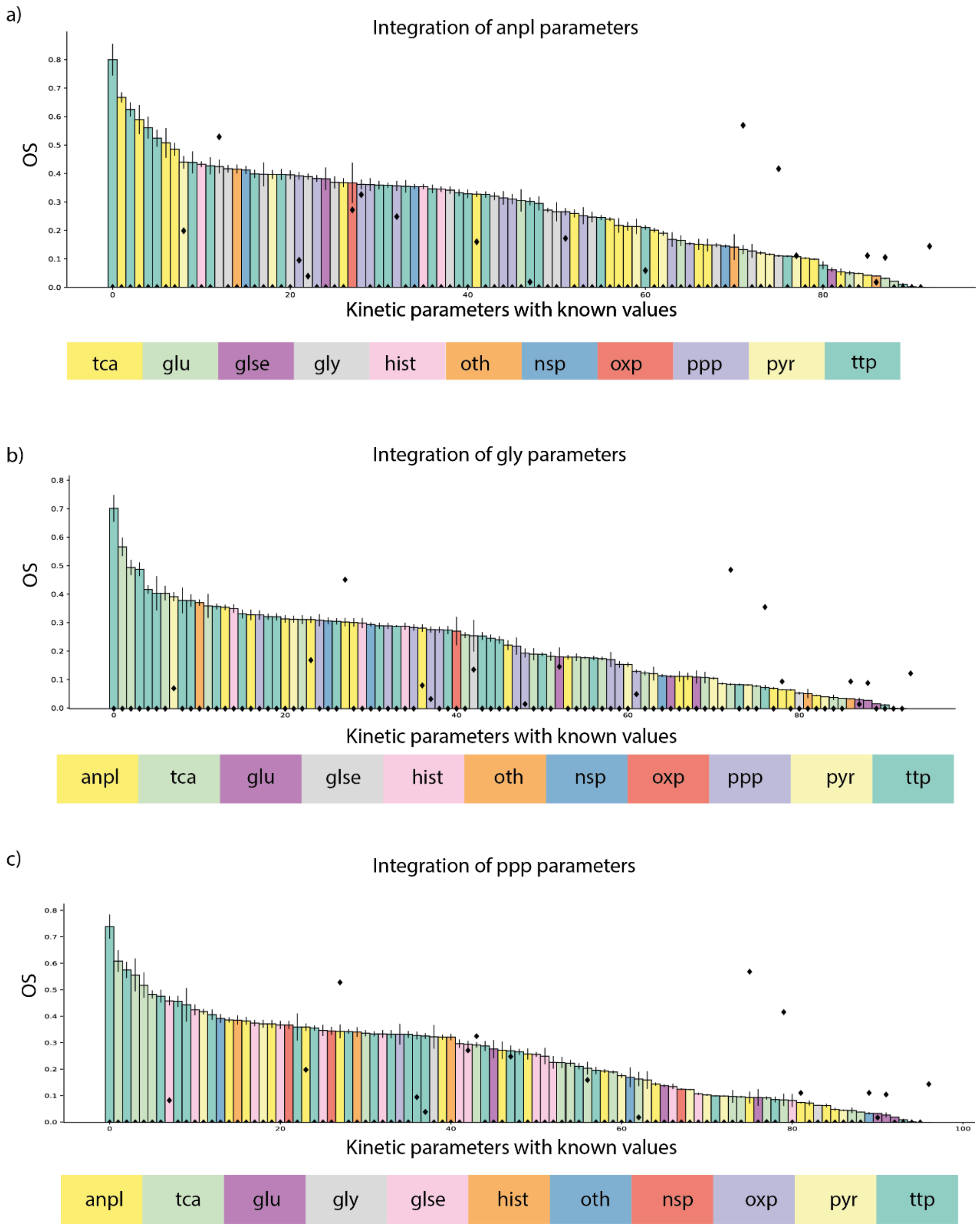
a)
Integration of anpl parameters
OS
Kinetic parameters with known values
tca
glu
glse
gly
hist
oth
nsp
oxp
ppp
pyr
ttp
b)
Integration of gly parameters
OS
Kinetic parameters with known values
anpl
tca
glu
glse
hist
oth
nsp
oxp
ppp
pyr
ttp
c)
Integration of ppp parameters
OS
Kinetic parameters with known values
anpl
tca
glu
gly
glse
hist
oth
nsp
oxp
pyr
ttp

d)

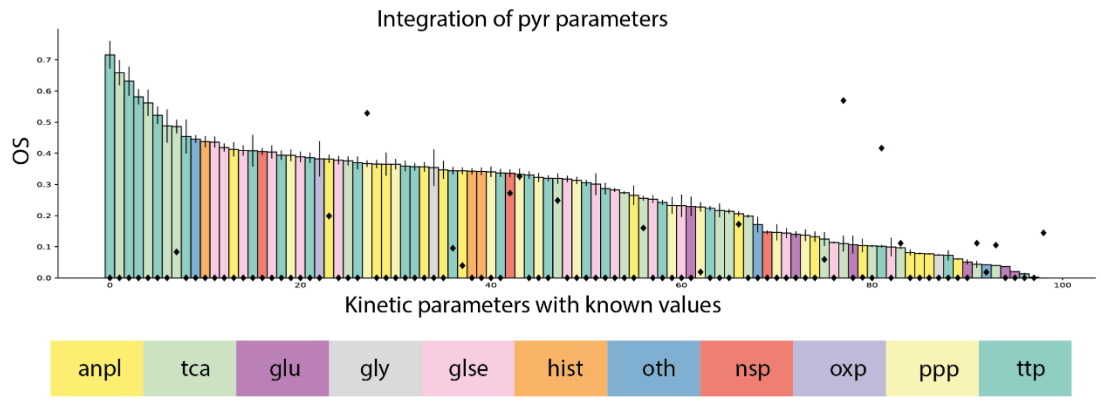


e)

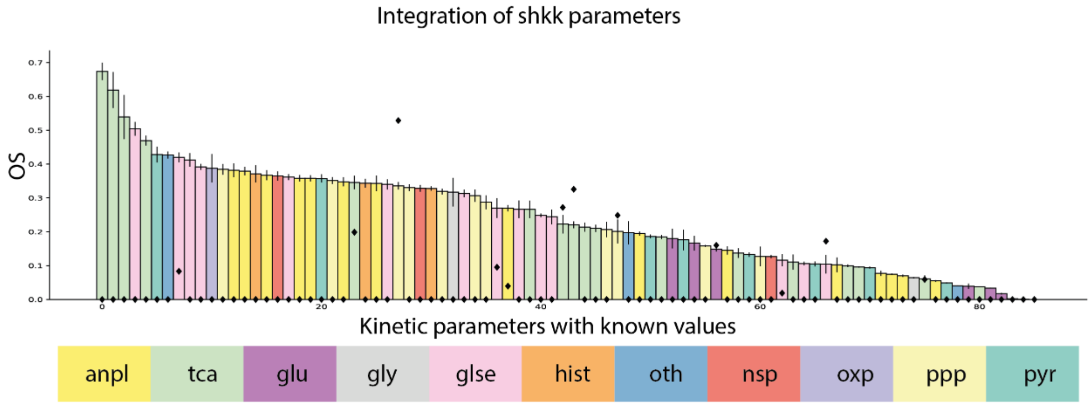


**Supplementary Figure. 10:** The mean overlap scores (OS) of all experimentally known $\boldsymbol{K_M}$s when $\boldsymbol{K_M}$s belonging to **(a)** anpl **(b)** gly **(c)** ppp **(d)** pyr and **(e)** shkk are integrated into RENAISSANCE (colored bars). The error bars represent the standard error in the OS. The black diamonds represent the OS when no $\boldsymbol{K_M}$s are integrated. Abbreviations: **ppp:** Pentose Phosphate Pathway, **gly:** Glycolysis/Gluconucleogenesis, **anpl:** Anaplerotic reactions, **shkk:** Shikimate pathway, **pyr:** Pyruvate metabolism.
